# Combinations of Indole based alkaloids from *Mitragyna speciosa* (Kratom) and cisplatin inhibit cell proliferation and migration of Nasopharyngeal carcinoma cell lines

**DOI:** 10.1101/2020.12.18.423377

**Authors:** Gregory Domnic, Nelson Jeng-Yeou Chear, Siti Fairus Abdul Rahman, Surash Ramanathan, Kwok-Wai Lo, Darshan Singh, Nethia Mohana-Kumaran

## Abstract

**Ethnopharmacological relevance:** *Mitragyna speciosa* (Korth.) or kratom is a medicinal plant indigenous to Southeast Asia. The leaves of *M. speciosa* is used as a medication in pain management including cancer related pain, in a similar way as opioids and cannabis. Despite its well-known analgesic effect, there is a scarce of information on the cancer-suppressing potential of *M. speciosa* and its active constituents.

**Aim of the study:** To assess the potential applicability of *M. speciosa* alkaloids (mitragynine, speciociliatine or paynantheine) as chemosensitizers for cisplatin in Nasopharyngeal carcinoma (NPC) cell lines.

**Materials and Methods:** The cytotoxic effects of the extracts, fractions and compounds were determined by conducting *in vitro* cytotoxicity assays. Based on the cytotoxic screening, the alkaloid extract of *M. speciosa* exhibited potent inhibitory effect on the NPC cell line HK-1, and therefore, was chosen for further fractionation and purification. NPC cell lines HK-1 and C666-1 were treated with combinations of cisplatin and *M. speciosa* alkaloids in 2D monolayer culture. The effect of the drug combination on cell migration was tested using *in vitro* wound healing and spheroid invasion assays.

**Results:** In our bioassay guided isolation, both methanolic and alkaloid extracts showed mild to moderate cytotoxic effect against the HK-1 cell line. Both NPC cell lines were insensitive to single agent and combination treatments of the *M. speciosa* alkaloids. However, mitragynine and speciociliatine sensitised the HK-1 and C666-1 to cisplatin ~4- and >5-fold, respectively in 2D monolayer culture. The combination of mitragynine and cisplatin also significantly inhibited cell migration of the NPC cell lines. Similarly, combination of mitragynine and cisplatin inhibited growth and invasion of HK-1 spheroids in a dose-dependent manner. Moreover, the spheroids did not rapidly develop resistance to the drug combinations at high concentrations over 10 days.

**Conclusion:** Collectively, data shows that both mitragynine and speciociliatine could be potential chemosensitizers for cisplatin. Further extensive drug mechanistic studies and investigations in animal models are necessary to delineate the applicability of *M. speciosa* alkaloids for NPC therapy.

## 1. Introduction

Nasopharyngeal carcinoma (NPC) is a malignant cancer which manifests in the nasopharynx, specifically in the pharyngeal recess, also known as the Fossa of Rosenmüller (Wei and Sham, 2005). The cancer is endemic in world regions namely South-Eastern Asia, Micronesia and Eastern Asia (Globocan, 2020). Concurrent cisplatin-based chemoradiotherapy is the standard of care treatment for NPC. Patients usually respond well in the early stages of cisplatin treatment, but gradually develop resistance to the drug, severely limiting its use in subsequent treatment episodes (Köberle et al., 2010). Combining cisplatin with compounds purified from natural resources is seen as a promising option to resensitize cancer cells to cisplatin. Indeed, compounds from natural resources have shown promise as an alternative treatment option given its reduced toxicity risk, affordability, and usage as a good treatment sensitizer (Lin et al., 2020).

*Mitragyna speciosa* (Korth.) or kratom is a medicinal plant, indigenous to Southeast Asia (Domnic et al., 2021). Historically, kratom leaves have been used as a remedy for treating common ailments such as fever, cough, pain, cancer, as a stimulant drug and opiate substitute (Saingam et al., 2012; Singh et al., 2016; Domnic et al., 2021). *M. speciosa* is rich in monoterpene indole and oxindole alkaloids, in which more than 40 alkaloids have been discovered (Brown et al., 2017). The pain-relieving action of *M. speciosa* is mainly attributed to its major opioid-like MIAs – mitragynine and speciociliatine (Sharma et al., 2019; Obeng et al., 2020). These alkaloids exert central analgesic activity through the agonistic action on μ-opioid receptor (Kruegel at al., 2016; Obeng et al., 2020). In traditional context, kratom is commonly used to treat cancer (Saingam et al., 2012). However, its widespread applicability as an alternative to cancer treatment remains inadequately documented. A previous *in vitro* study showed that cancer cell lines were sensitive to mitragynine at higher concentrations (Goh et al., 2014). Another study reported that mitragynine exerted dose-dependent cytotoxic effects on human cancer cells, SH-SY5Y and HEK 293 (Saidin et al., 2008).

Given that kratom is anecdotally used to treat cancer, and due to the lack of studies on its therapeutic properties, thus, we aim to investigate the sensitivity of NPC cancer cell lines to combination of *M. speciosa* alkaloids and cisplatin, in 2D monolayer culture and in 3-dimensional spheroids, which resemble a microenvironment and architecture closer to tumours *in vivo* (Siva Sankar et al., 2017).

## 2. Materials & methods

### 2.1 Plant material

Fresh leaves of *Mitragyna speciosa* (Korth.) (Rubiaceae) were collected from a kratom plantation in Permatang Rawa, Penang, Malaysia (5°22’17.0″N 100°27’01.0″E). The taxonomic identity of the plant was authenticated by Dr. Rosazlina Rusly, a botanist from School of Biological Sciences, Universiti Sains Malaysia. A voucher specimen (NEL-(K)-2019(01)) was deposited at the Herbarium Unit of the premise.

### 2.2 Extraction of plant material

Powdered dry leaves of *M. speciosa* (0.5 kg) was extracted using hot maceration method with 5 L of methanol at 45 °C for 3 consecutive days according to the procedures described (Chear et al., 2021). The suspension was filtered, and the supernatant was evaporated to dryness at 40 °C under reduced pressure to afford 150.0 g of crude extract. The methanol extract was then re-suspended in 10% (*v/v*) acetic acid and stir for 24 h. The suspension was filtered, and the acidic solution was defatted with n-hexane (3 times). The defatted acidic solution was adjusted to pH 9-10 with 25% (*v/v*) ammonium hydroxide. The alkaline solution was then partitioned with chloroform to liberate the alkaloid constituents (free base). The collected chloroform fraction was concentrated *in vacuo* to afford alkaloid-enriched extract (5.5 g).

### 2.3 Bioassay guided isolation of M.speciosa alkaloids

Based on the cytotoxic screening, the alkaloid extract of *M. speciosa* exhibited potent inhibitory effect on NPC cell line HK-1 and therefore, was chosen for further fractionation and purification. Briefly, 5.5 g of alkaloid extract was fractionated on a silica gel column chromatography (CC) with a step gradient elution of hexane - ethyl acetate – methanol (100:0:0 to 0:0:100, v/v) to produce 7 major fractions – F1 (0.05 g), F2 (2.12 g), F3 (0.56 g), F4 (0.43 g), F5 (0.31 g), F6 (0.82 g) and F7 (0.21 g). Both active F2 (% inhibition: 53% at 100 μg/mL) and F6 (% inhibition: 74% at 100 μg/mL) were subjected to compound purification. Approximately 1.0 g of sample was loaded on a medium size silica gel CC (5 x 40 cm) and eluted isocratically with a mixture of hexane and ethyl acetate (80:20, *v/v*) to afford mitragynine (0.76 g) (**1**) for F2. F6 (0.5 g) was further purified on a medium size silica gel CC (5 x 40 cm) and eluted with a step gradient of ethyl acetate-methanol to afford speciociliatine (0.12 g) (**2**). Other major alkaloid standards – paynantheine (**3**) and speciogynine (**4**) were purified from F3 and F4, respectively by a silica gel flash chromatography (hexane-ethyl acetate, 90:10 to 20:80, *v/v*) (Chear et al., 2021). The chemical identity of each isolated compound was confirmed with the aid of ^1^H & ^13^C NMR, MS, and the spectroscopic data was then compared with those published data in literature (Sharma et al., 2019; Obeng et al., 2020). The purity level of each alkaloid was evaluated by HPLC-PDA according to the method described by (Saref et al., 2019).

### 2.4. Compounds and cell lines

The *M. speciosa* alkaloids were dissolved in dimethyl sulfoxide (DMSO) at a stock concentration of 10 mM. The HK-1 and C666-1 cell lines were cultured in RPMI 1640 medium supplemented with 10% heated foetal bovine serum (FBS). Additionally, 10% (v/v) Glutamax was added to the RPMI complete medium that was used to grow the C666-1 cells. NPC cell lines HK-1 and C666-1 were authenticated using the AmpFISTR profiling as described previously (Daker et al., 2012).

### 2.5 M. speciosa extracts and fractions cytotoxicity assay

The cytotoxic effects of the extracts and fractions were determined by conducting *in vitro* cytotoxicity assays, and were performed as previously described (Allen et al., 2002). Initially, 300 μg/ml of the crude extract, 100 μg/ml of the alkaloid extract and 25 μg/ml of each column fraction (1-7) was tested against the HK-1 cell line and, from the results, the active fractions were considered to be those which gave less than 50% survival at an exposure time of 72 h. From the initial screening of the extracts and fractions, the crude extract, alkaloid extract and column fractions (1-7) were further diluted (two-fold steps) in medium to produce 8 concentrations respectively (Crude extract = 2.34, 4.69, 9.38, 18.75, 37.5, 75, 150 and 300 μg/ml; Alkaloid extract = 0.78, 1.56, 3.13, 6.25, 12.5, 25, 50 and 100 μg/ml; Column fractions (1-7) = 0.39, 0.78, 1.56, 3.13, 6.25, 12.5, 25 and 50 μg/ml). One hundred μl/well of each concentration of drugs was added to the plates in four replicates. The final dilution used for treating the cells contained not more than 1% of the initial solvent, this concentration being used in the solvent control wells. The plates were incubated for 72 h.

### 2.6 Combination of M. speciosa alkaloids and cisplatin cytotoxicity assays

Drug sensitivity assays were performed as described previously (Allen et al., 2002). NPC cell lines HK-1 and C666-1 were treated with either *M. speciosa* compounds; mitragynine (1), speciociliatine (2), paynantheine (3) or speciogynine (4) diluted in two-fold steps (0.25, 0.5, 1, 2, 4, 8, 16 and 32 μM) either alone or in combination for 72 h.

Sensitivity of the HK-1 cells to drug combinations was measured by testing a fixed concentration of either mitragynine, speciociliatine or paynantheine at 8, 16 or 32 μM with increasing concentrations of cisplatin diluted in two-fold steps (0.25, 0.5, 1, 2, 4, 8, 16 and 32 μM). Cisplatin (MedChemExpress, NJ, USA) was initially prepared by dissolving the powder in 0.9% salt solution at a stock concentration of 5 mM. The C666-1 cells were treated similarly except that a fixed concentration of either mitragynine, speciociliatine or paynantheine at 25, 35 or 45 μM was employed. Cell proliferation was quantified by fluorescence using SYBR Green as described previously (Lian et al., 2018). The drug sensitivity assays were performed four times (n=4) and mean IC_50_ values were calculated from the experimental data. The dose curves were generated as described previously (Abdul Rahman et al., 2020). The x-axis was formatted to have a scale of base 10 logarithm, but the drug concentrations employed were not log-transformed prior to plotting the graphs. The CalcuSyn 2.11 program (Biosoft, Cambridge, UK) was used to access drug interactions.

### 2.7 Wound healing assay

The NPC cell lines C666-1 and HK-1 cells were seeded at a density of 5 x10^5^ and 2 x10^5^ cells/well, respectively in complete medium and grown overnight to a 90% confluent monolayer, in 24-well culture plates. After incubation, scratch wounds were generated using a sterile 200 μl pipette tip by pressing firmly against the top of the tissue culture plate and swiftly making a vertical wound down through the cell monolayer. The detached cells and debris were removed by gentle washing with 500 μL PBS. The HK-1 and C666-1 cells were treated with cisplatin and mitragynine alone and in combination for 48/72 h, respectively. The concentrations of mitragynine and cisplatin (single treatments and combination) used for the assay was based on the IC_50_ values obtained from the cytotoxicity assays. The cells were treated with media supplemented with 5% foetal bovine serum to prevent cell proliferation from interfering with cell migration. Images of cell migration (three fields of each triplicate well) were captured using an inverted phase-contrast microscope (Olympus CKX41; Olympus, Tokyo, Japan) at 48/72 h time interval for the HK-1 and C666-1, respectively. The numbers of cells that migrated into the wound area were evaluated using the formula: Percentage of migrated cells = [initial scratch (0 h) - final scratch (48 h/72 h)]/initial (0 h) x 100. The analyses of the cell migration images were performed using the Fiji software (v1.51s, NIH) (Schindelin et al., 2012) which calculates the scratch area (= open wound area) for each image.

### 2.8 Three-dimensional spheroids

Spheroids were generated from the HK-1 cells and embedded in collagen matrix as previously described (Smalley et al., 2008; Abdul Rahman et al., 2020). Spheroids were treated with cisplatin and mitragynine alone and in combination. Drug/compound and media were replenished every 72 hours for experiments conducted over 10 days. Spheroids growth and invasion were photographed using the Nikon C2+ inverted fluorescence microscope. Images were processed using ImageJ (v1.51s, NIH) (Schindelin et al., 2012).

### 2.9 Statistical analysis

Data was represented as mean ± SEM of three experiments for the wound healing assay. Independent sample t-tests and one-way ANOVA were used to compare the statistical differences between the groups. P values <0.05 was considered statistically significant.

## 3. Results

### 3.1 Bioassay-guided isolation of cytotoxic compounds against NPC cell lines

According to the general screening protocol, cytotoxicity of plant extracts was scored into four categories namely; very active (IC_50_◻≤◻20 μg/mL), moderately active (IC_50_◻>◻20–100 μg/mL), weakly active (IC_50_◻>◻100–1000 μg/mL) and inactive (IC_50_◻>◻1000 μg/mL) (Atjanasuppat et al., 2009; Baharum et al., 2014). In this study, the methanolic leaf extract of *M. speciosa* and its alkaloid enriched extract were tested for their potential cytotoxic effect on HK-1 cell line using the Sybr Green I assay. Both methanolic and alkaloid extracts showed mild to moderate cytotoxic effect against HK-1 cell line with IC_50_ values of 133.71 and 32.16 μg/mL, respectively (Table 1). The cytotoxicity of alkaloid extract was approximately 5 times greater than its origin – methanolic extract. Therefore, the alkaloid extract was considered as an active extract and regarded for further evaluation for its cytotoxic constituents. Alkaloid extract was further fractionated to yield seven major fractions, F1-F7. Among them, only F2 and F6 exhibited strong inhibitory activity against HK-1 cells with IC_50_ values of 19.65 and 11.55 μg/mL, respectively (Table 1). Fractions F2 and F6 were then purified on silica gel column chromatography to afford two indole- based *M. speciosa* alkaloids - mitragynine (**1**) and speciociliatine (**2**), respectively. In order to understand the structure-cytotoxicity relationship of *M. speciosa* alkaloids, two other mitragynine derivatives or diastereoisomers - paynantheine (**3**) and speciogynine (**4**) were also isolated from F3 and F4, respectively. ^1^H & ^13^C NMR and MS spectroscopic analysis (Supplementary figures 1-16) showed that compound **1**, **2**, **3** and **4** presented analytical data in full agreement with the published data in the literature (Sharma et al., 2019; Obeng et al., 2020). The chemical structures of identified compounds (**1**-**4**) are shown in Fig. 1.

**Fig. 1:**
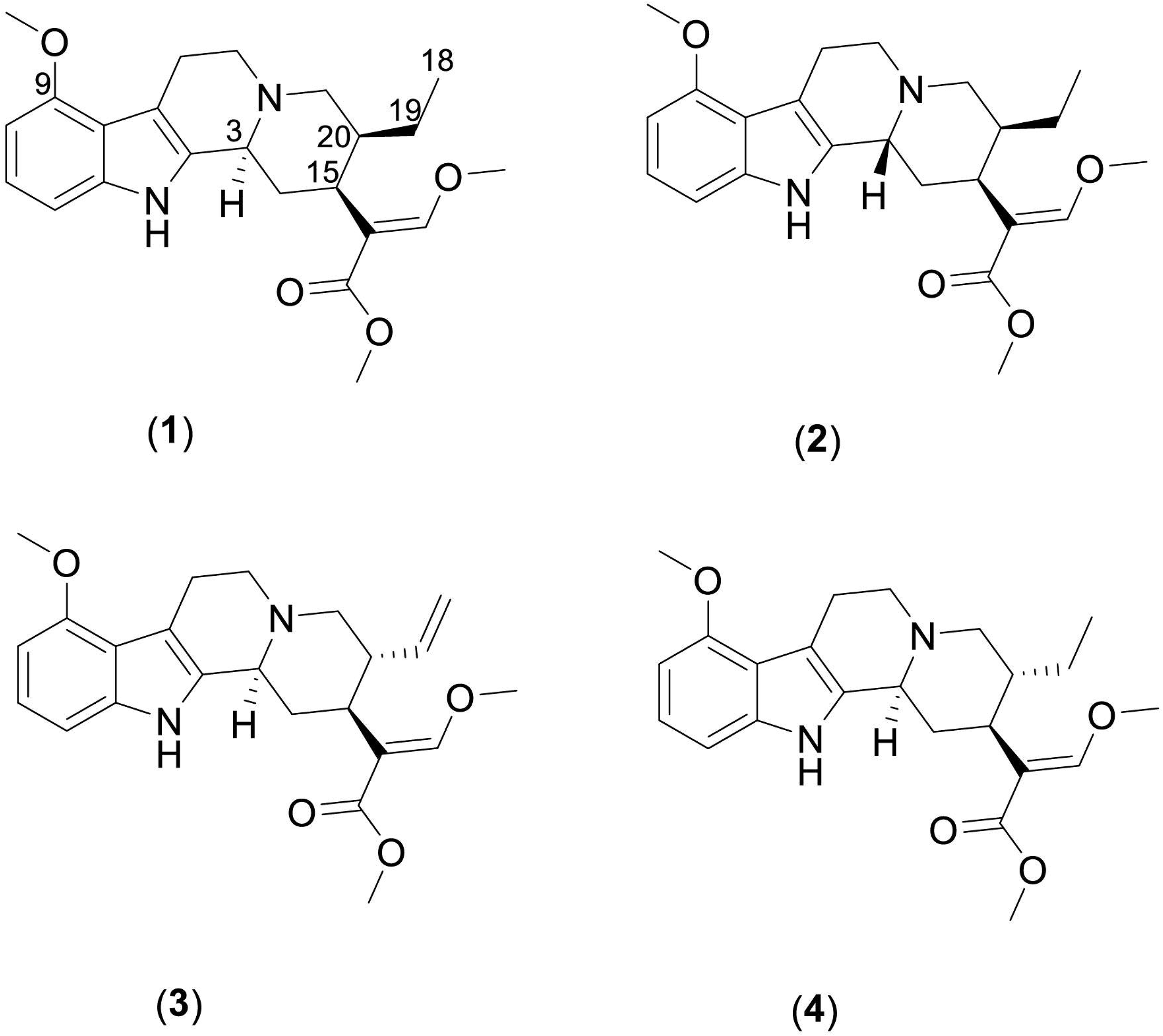
Chemical structures of the isolated *M.speciosa* alkaloids (mitragynine (**1**), speciociliatine (**2**), paynantheine (**3**) and speciogynine (**4**).

### 3.2 NPC cell lines were insensitive to single agent treatment of M. speciosa alkaloids

The NPC cell lines HK-1 and C666-1 were subjected to single agent treatment of all four *M. speciosa* alkaloids. Both cell lines were resistant to single agent treatment of *M. speciosa* alkaloids (**1**-**4**) except at high treatment doses (Supplementary Table 1). Given that the cell lines were resistant to single agent treatment of the *M. speciosa* alkaloids, the HK-1 cells were subjected to combination of mitragynine and speciociliatine at a 1:1 compound concentration ratio. The cells were insensitive to the combination (IC_50_: 26.10 ± 1.21 μM; Supplementary table 2). In order to test the sensitization effects of the compounds, a fixed dose of an alkaloid was added to escalating doses of another. The HK-1 cells were treated with increasing concentrations of mitragynine and 10 μM speciociliatine and *vice versa* (10 μM was the sublethal dose of the alkaloids in the single agent treatment). However, in all the compound combination scenarios presented, the cells exhibited insensitive to the compounds (Supplementary Table 2). Compound interaction analyses indicated that none of the combination tested were synergistic (Supplementary Table 2).

### 3.3 Sensitization of NPC cells to cisplatin by M. speciosa alkaloids

Given that the NPC cell lines were insensitive to single agent treatment of the *M. speciosa* alkaloids, we investigated the potential of these compounds as chemosensitizers for cisplatin. NPC cells were treated with a fixed concentration of either mitragynine, speciociliatine or paynantheine with increasing concentrations of cisplatin.

The HK-1 and the C666-1 cells were resistant to single agent treatment of cisplatin and *M. speciosa* compounds (Fig. 2A and Supplementary table 3). The HK-1 cells were sensitized to cisplatin by 2.2-fold, at 8 μM of mitragynine. The sensitization increased to 3.5-fold and 4-fold at a concentration of 16 μM and 32 μM mitragynine (Fig. 2A and Table 2). In C666-1 cells, mitragynine at 25 μM and 35 μM only sensitized C666-1 cells to mitragynine by 2-fold. The sensitization increased to > 5-fold when the concentration of mitragynine was increased to 45 μM (Fig. 2B and Table 3).

**Fig. 2:**
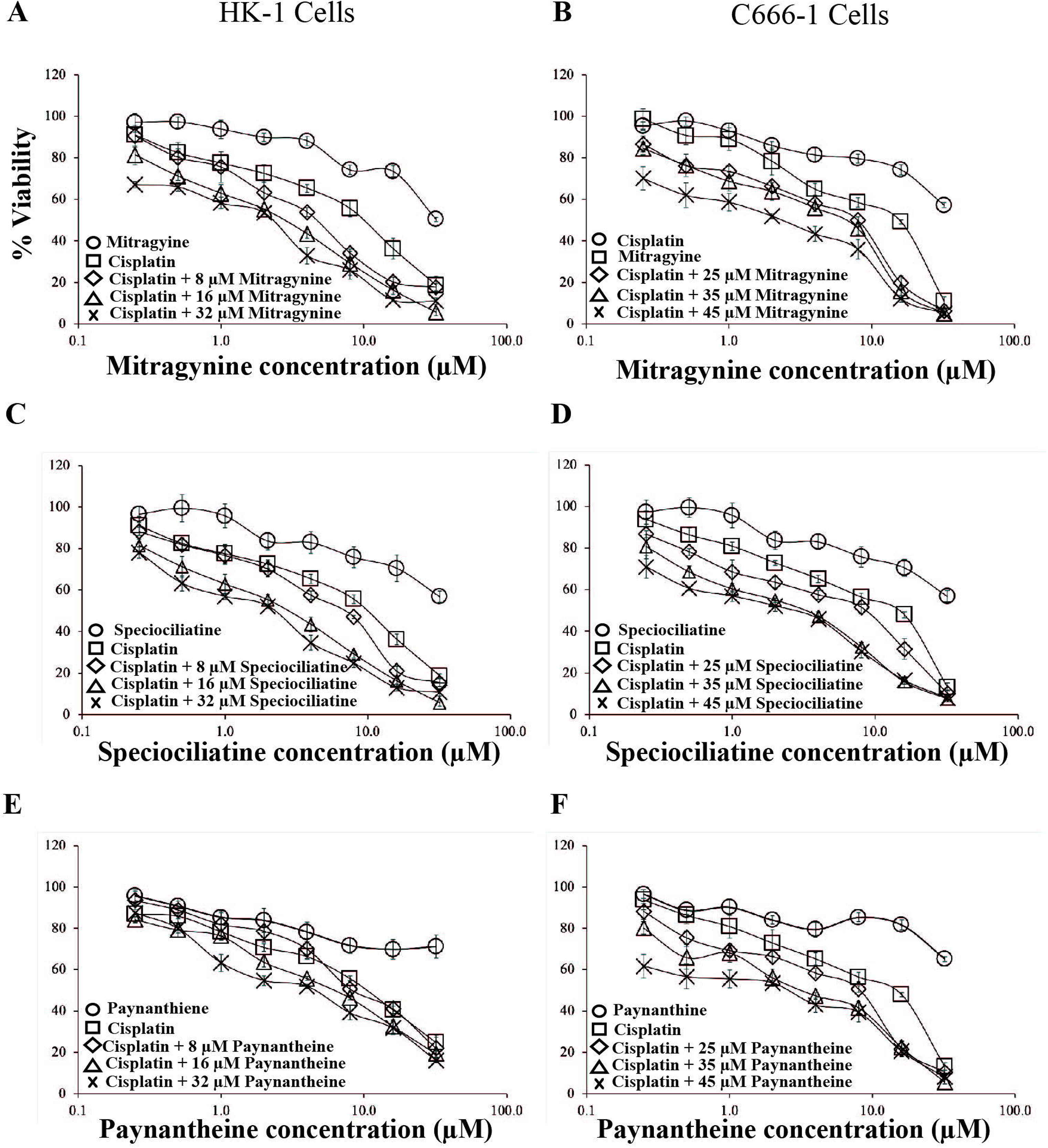
Sensitization of the NPC cell lines to cisplatin by the mitragyna alkaloids. NPC cell lines HK-1 and C666-1 were treated with increasing concentrations of cisplatin (0-32 μM) in the presence and absence of **(A-B)** Mitragynine; **(C-D)** Speciociliatine and **(E-F)** Paynanthiene. Sensitivity of the NPC cell lines to single agent treatment of either Mitragynine, Speciociliatine or Paynantheine are also shown in each dose-response graph. Points represent mean ± SEM of four experiments.

Combination with 8 μM of speciociliatine sensitized the HK-1 cells to cisplatin by ~ 2-fold. The sensitization increased to 3-fold and 4-fold when the concentration of speciociliatine was increased to 16 μM and 32 μM (Fig. 2C and Table 2), respectively. In C666-1 cells, speciociliatine at 25 μM sensitized the C666-1 cells to cisplatin by ~ 2-fold. The sensitization increased by 4-fold and by > 5-fold at 35 μM and 45 μM (Fig. 2D and Table 3), respectively.

In HK-1 cells, the presence of either 8 μM or 16 μM or 32 μM paynantheine, sensitized the cells to cisplatin only by ~ 2-fold (Fig. 2E and Table 2). In C666-1 cells paynantheine at 25 μM sensitized the C666-1 cells to cisplatin by ~ 2-fold. The sensitization increased to by 4-fold and > 5-fold when the concentration of paynantheine was increased to 35 μM and 45 μM (Fig. 2F and Table 3), respectively.

Drug interaction analyses indicated that combination of the *M. speciosa* alkaloids (mitragynine, speciociliatine or paynantheine) and cisplatin demonstrated synergism at several concentrations in both cell lines (Figs. 3 and 4).

**Fig 3:**
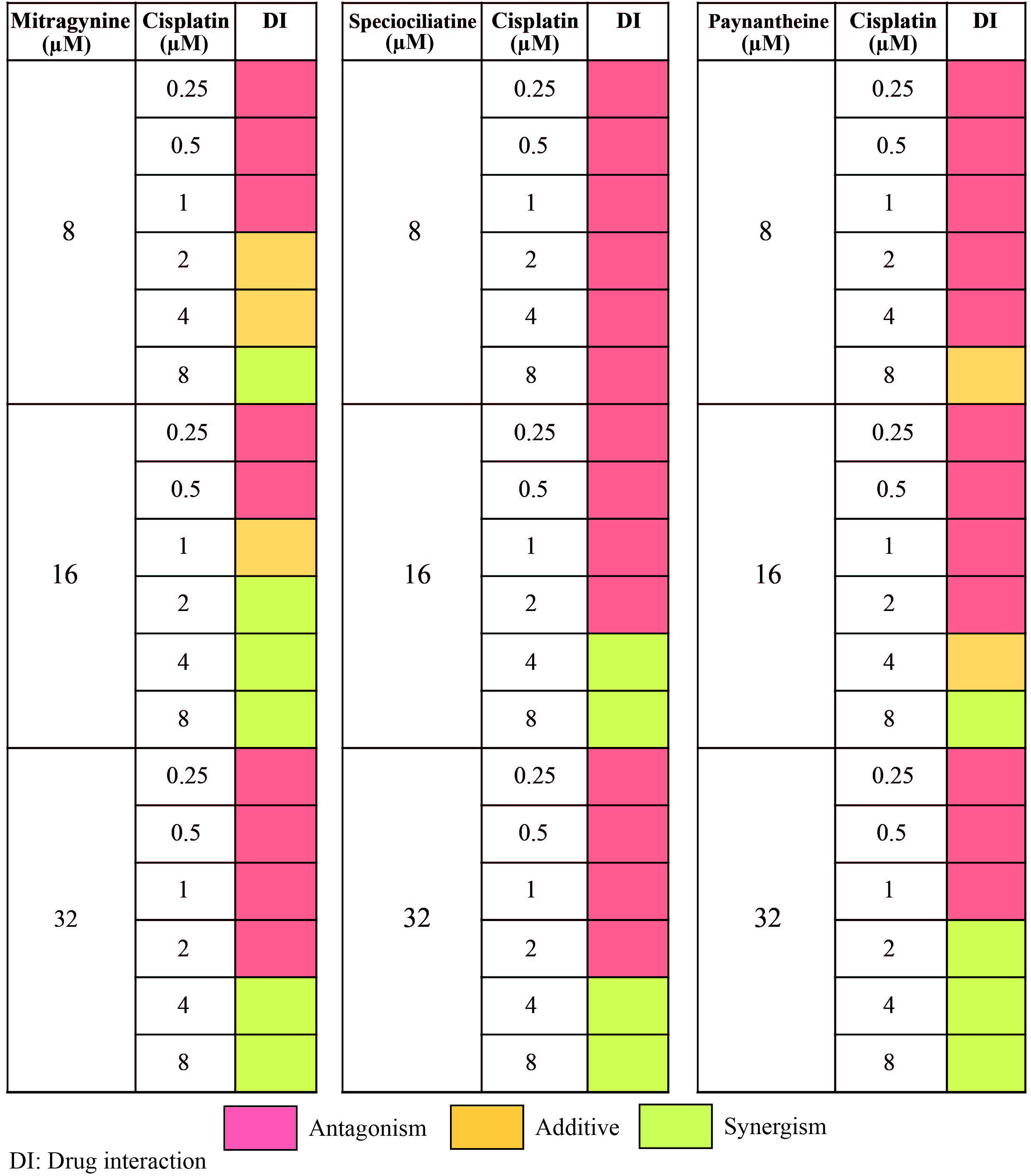
Drug combination interactions of sensitization of the HK-1 cells to cisplatin by mitragynine. The drug interactions were determined from the combination index (CI) values generated from the CalcuSyn 2.11 software (Biosoft, Cambridge, UK).

**Fig 4:**
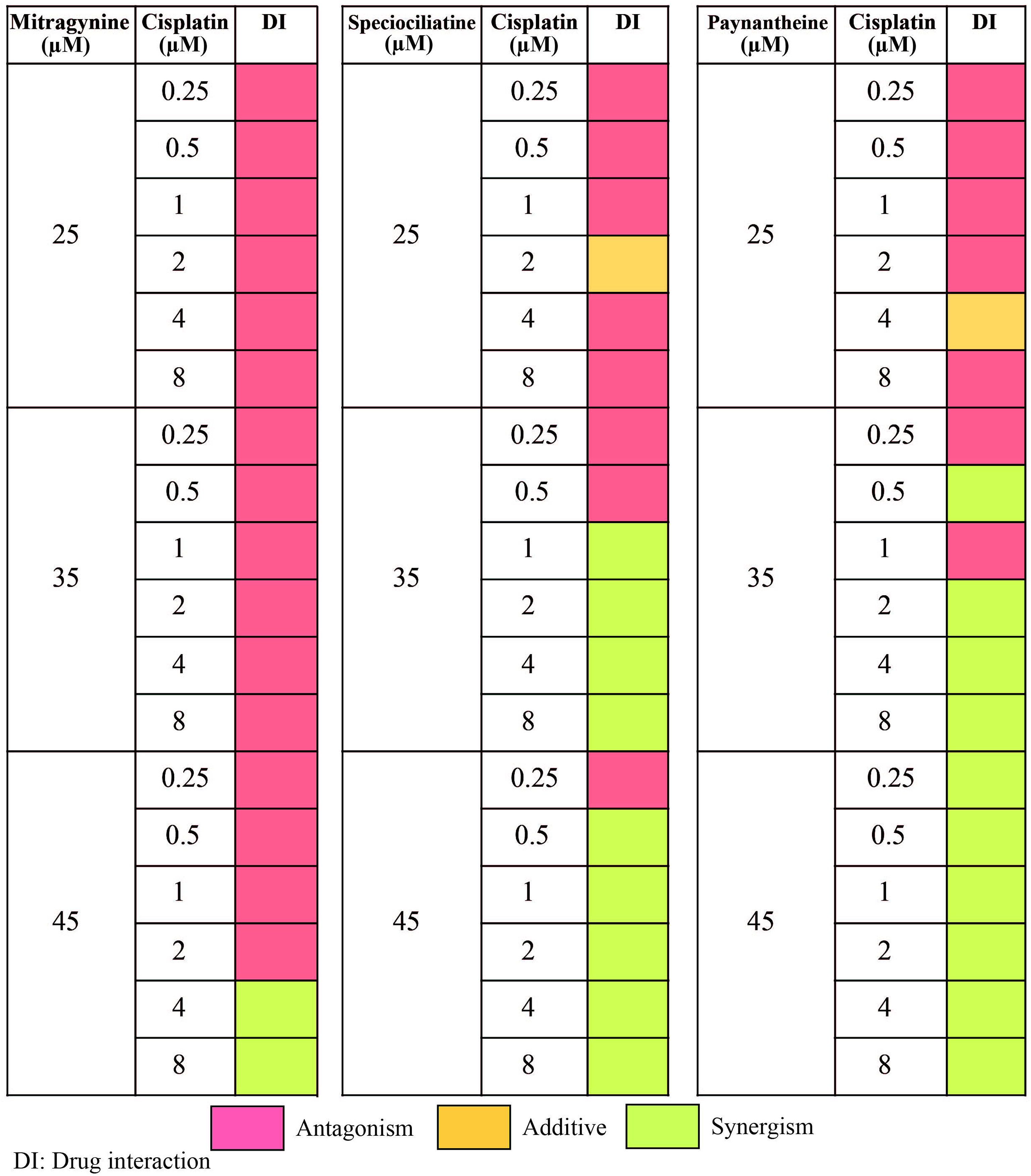
Drug combination interactions of sensitization of the C666-1 cells to cisplatin by mitragynine. The drug interactions were determined from the combination index (CI) values generated from the CalcuSyn 2.11 software (Biosoft, Cambridge, UK).

### 3.4 Combination of mitragynine and cisplatin inhibit growth and invasion of HK-1 cells

In order to evaluate the effects of mitragynine either alone or in combination with cisplatin to inhibit the migration of the NPC cell line HK-1 and C666-1; wound healing assay was performed. Treatment induced effect on wound closure was computed by measuring the differences in area of wounds at 0 h (immediately after wounding) and at different end time points. The serum concentration throughout the drug treatment groups and the controls were kept low (RPMI + 5% FBS), to prevent cells proliferation. A higher cell migration was noted for the cisplatin treatment alone in the HK-1 cells (42%), while an opposite effect was noted in the C666-1 cells (11%). In the mitragynine-treated group, it was noted that there was no significant inhibition of migration in both the HK-1 (80%) and C666-1 (25%) cells. The combination of mitragynine and cisplatin, however, showed a reduction in the percentage of cell migration from 80% (mitragynine) to 11% (combined) in the HK-1 cells (Fig. 5A), and similarly from 25 (mitragynine) to 4% (combined) in the C666-1 cells (Fig. 5B). The assay demonstrated the potential of the combination of mitragynine and cisplatin in significantly inhibiting the migration across both cell lines as compared with the control and single agent groups (mitragynine and cisplatin).

**Fig 5:**
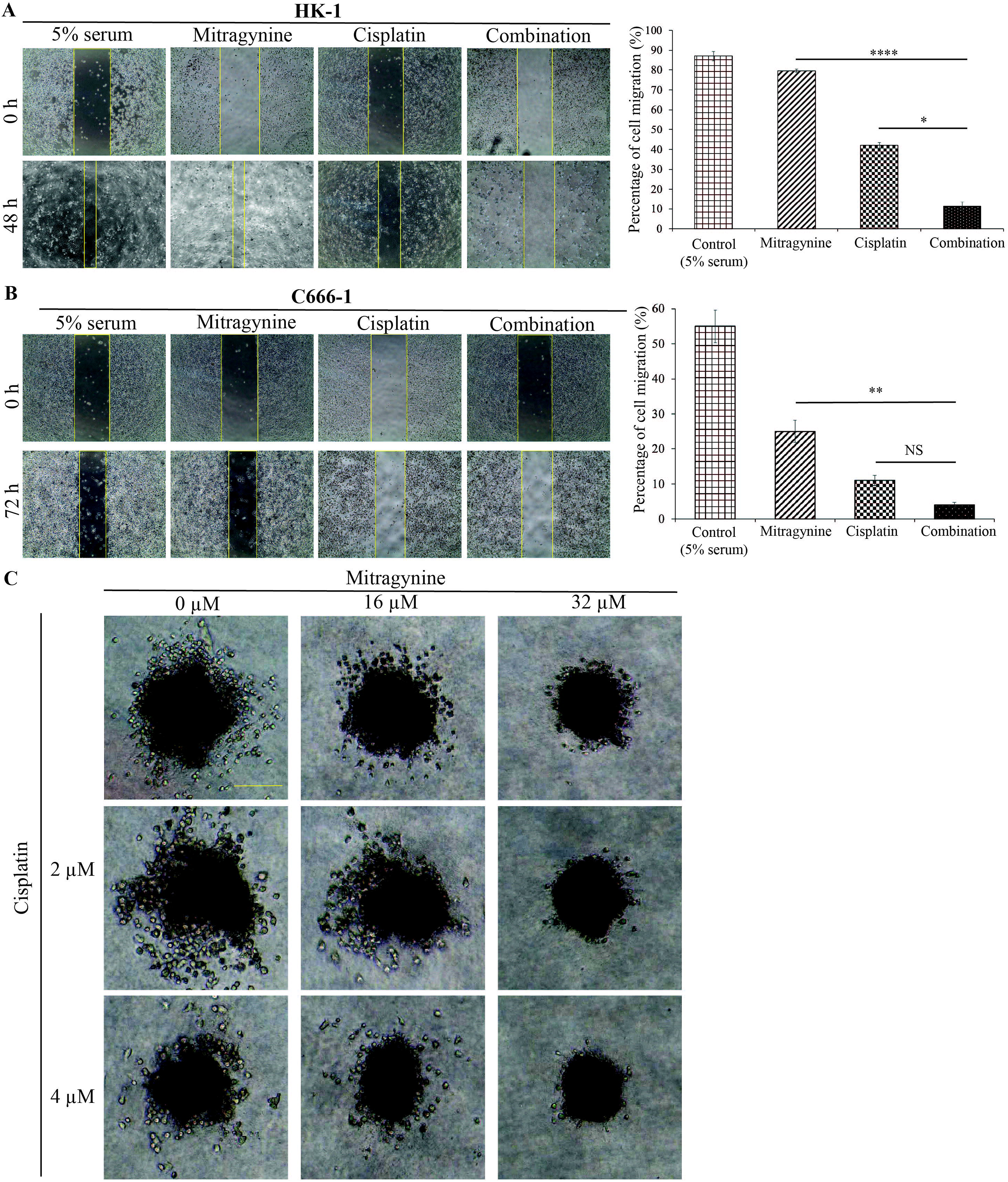
Combination of mitragynine and cisplatin inhibits NPC cell migration and invasion. Representative images and analysis of wound closure are shown for **(A)** HK-1 after 48 h and **(B)** C666-1 after 72 h of mitragynine and cisplatin treatment alone and in combination using the IC_50_ values (Fig 2. A-B). Bar graphs show percentage differences in wound closure after 48 h in the HK-1 cells and 72 h in the C666-1 cells. Error bars indicate standard error of mean. Statistically significant differences in wound closure between the treatment groups are shown as ****p < 0.0001 or **p < 0.01 or *p < 0.05 determined by two-tailed paired T-test. **(C)** The effect of combination of mitragynine and cisplatin on the growth and invasion of HK-1 spheroids over three days. The spheroids were treated with single agents mitragynine and Cisplatin and combination of both over three days at the indicated concentrations, n = 2-3 spheroids per combination. Size bar: 200 μm.

Next, we utilized a 3D spheroid invasion assay as a more representative model of in vivo invasion. Given that the HK-1 cells form compact spheroids compared to the C666-1 cells (Lian et al., 2020), the HK-1 cells were used for the spheroid study. The effect of combination of cisplatin and mitragynine on the growth and invasion of the spheroids were first studied over 3 days of treatment. Combination of 16 μM of mitragynine and 4 μM of cisplatin reduced spheroid growth and invasion compared to single agent treatment of mitragynine and cisplatin (Fig. 5C).

The spheroids were monitored till day 10 for emergence of resistant cells at high concentrations of the combination. Inhibition of spheroid growth and invasion were already obvious at combination of 16 μM of mitragynine and 2 μM of cisplatin. The reduction in spheroid growth and invasion was more significant in the presence of 16 μM of mitragynine and 4 μM of cisplatin (Fig. 6). Moreover, the spheroids did not rapidly develop resistance to the drug combination at high concentrations.

**Fig 6:**
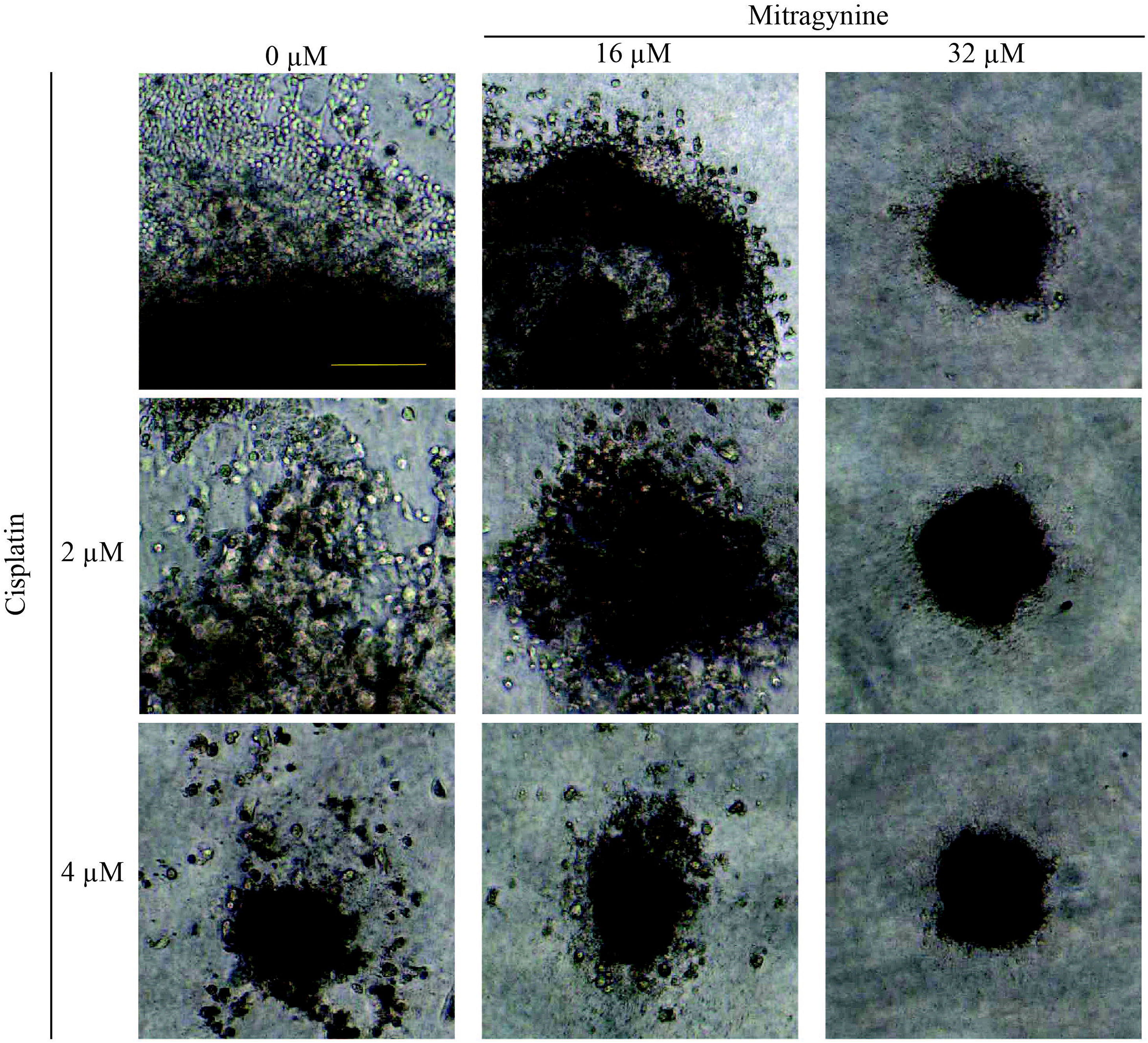
The effect of combination of mitragynine and cisplatin on the growth and invasion of HK-1 spheroids over ten days. The spheroids were treated with single agents mitragynine and cisplatin and combination of both over ten days at the indicated concentrations, n = 2-3 spheroids per combination. Media and compound/drug were replenished every 72 h. There were no resistance cells observed at the highest concentrations of the combination over 10 days. Size bar: 200 μm.

## 4. Discussion and Conclusion

Our findings highlight the medicinal utility of *M. speciosa* alkaloids as potential sensitizers for nasopharyngeal cancer treatment. In single agent treatment, all tested *M. speciosa* alkaloids – mitragynine **(1)**, speciociliatine **(2)**, paynantheine **(3)** and speciogynine **(4)** showed weak to moderate inhibition against nasopharyngeal carcinomas - HK-1 and C666-1 cell lines. Among these alkaloids, mitragynine **(1)** showed the highest inhibition against the growth of HK-1 cells, followed by speciociliatine **(2)**. Other tested alkaloids were found to be non-cytotoxic at highest dose. Based on the structure-activity relationships (SARs) study, *R* orientation at C-3 and S orientation at C-20 positions are the key features for the cytotoxicity of mitragynine. Speciociliatine was found to exhibit weaker cytotoxicity than mitragynine due to the change of orientation from R to S at C-3 position. Similarly, the cytotoxicity of speciogynine and paynantheine were abolished due to the R orientation at its C-20 position.

The NPC cell lines were first tested to single agent treatment of all three *M. speciosa* alkaloids (mitragynine, speciociliatine and paynantheine) and cisplatin. Both NPC cell lines were resistant to single agent treatment of all three *M. speciosa* alkaloids and cisplatin except at high doses. Similar observations were reported in another study which investigated the sensitivity of HCT 116 (colon carcinoma cell line) and K562 (leukaemia cell line) to single agent mitragynine. Both cell lines were sensitive to mitragynine at concentrations > 100 μM (Goh et al., 2014). The sensitivity of cancer cells to other *M. speciosa* alkaloids has to our knowledge not yet been reported in other studies.

Mitragynine and speciociliatine sensitized the NPC cells to cisplatin > 4-fold at concentrations close to its IC_50_ values. These compounds might thus be effective chemosensitizers to cisplatin. The C666-1 cells were more sensitive to combination of paynantheine and cisplatin compared to HK-1 indicating that the effect of paynantheine is more cell-type specific.

Notably, we also observed that the combination of cisplatin and mitragynine suppressed cell migration and invasion. In addition, the findings of the wound healing assay were consistent with our spheroid studies, as when mitragynine is combined with cisplatin – a significant reduction in spheroid growth and invasion was observed in a dose-dependent manner, suggesting that the combination may be effective *in vivo*. Taken together, the *M. speciosa* alkaloids sensitized the NPC cells to cisplatin. Furthermore, the combination of mitragynine and cisplatin impaired wound closure in the wound healing assay, as well as suppressed invasion of the spheroids.

These findings suggest that the *M. speciosa* alkaloids hold potential to be developed as future chemosensitizers for the treatment of NPC. Prospective studies must attempt to describe the mechanism of action, as well as testing the combination of *M. speciosa* alkaloids and cisplatin in other cancer types and *in vivo* models to support its potential applicability for cancer treatment.

## Abbreviations

NPC: Nasopharyngeal carcinoma
2D: 2-dimensional
FBS: Foetal Bovine Serum
PBS: Phosphate Buffer Saline
NMR: Nuclear Magnetic Resonance
GC-MS: Gas Chromatography-Mass Spectrometry
HPLC-PDA: High-Performance Liquid Chromatography–Photodiode Array

## Author contributions

**Gregory Domnic:** Methodology, Formal analysis, Writing – Original Draft; **Nelson Jeng Yeou Chear**: Methodology, Formal analysis; Writing – Original Draft; **Siti Fairus Abdul Rahman:** Methodology, Formal analysis; **Surash Ramanathan:** Resources; **Kwok-Wai Lo:** Resources; **Darshan Singh:** Resources, Funding acquisition, Writing – Review & Editing; **Nethia Mohana-Kumaran:** Conceptualization, Methodology, Formal analysis, Resources, Writing – Original Draft, Funding acquisition.

## Declaration of competing interest

The authors declare that there are no conflicts of interest

## Acknowledgements

We would like to thank Professor Dr. George Sai Wah Tsao (University of Hong Kong, Pokfulam, Hong Kong, China) for providing the NPC cell line HK-1. DG is a recipient of the Biasiswa Yang Di-Pertuan Agong 2018/19. NJYC is a recipient of the Universiti Sains Malaysia Fellowship Scheme. This study was funded by the Universiti Sains Malaysia Research University (RU) grant (Grant number: 1001/PBIOLOGI/8012268) and the Fundamental Research Grant Scheme (FRGS), Ministry of Education, Malaysia (Grant number: 203/PBIOLOGI/601228).

## Notes

### Competing Interest Statement

The authors have declared no competing interest.

### Summary of Updates

Title revised Abstract revised Introduction revised Some results sections were revised and some were newly added. Discussion revised New images and supplementary images added.

