## Supplementary data for "Combinations of Indole based alkaloids from *Mitragyna speciosa* (Kratom) and cisplatin inhibit cell proliferation and migration of Nasopharyngeal carcinoma cell lines"

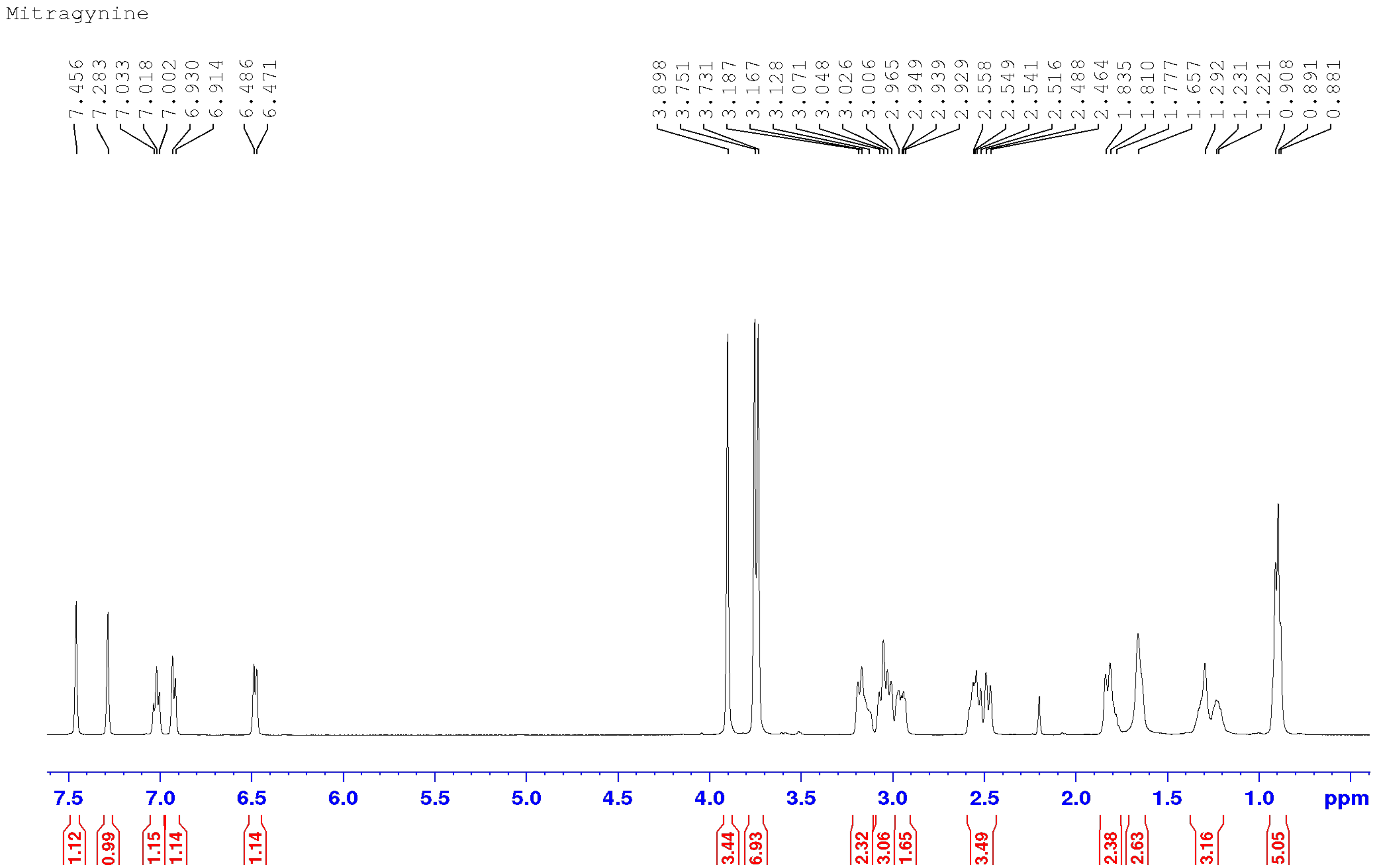


**Supplementary figure 1**: [1]H NMR spectrum of mitragynine (**1**) ($CDCL_3$, 500 MHz).

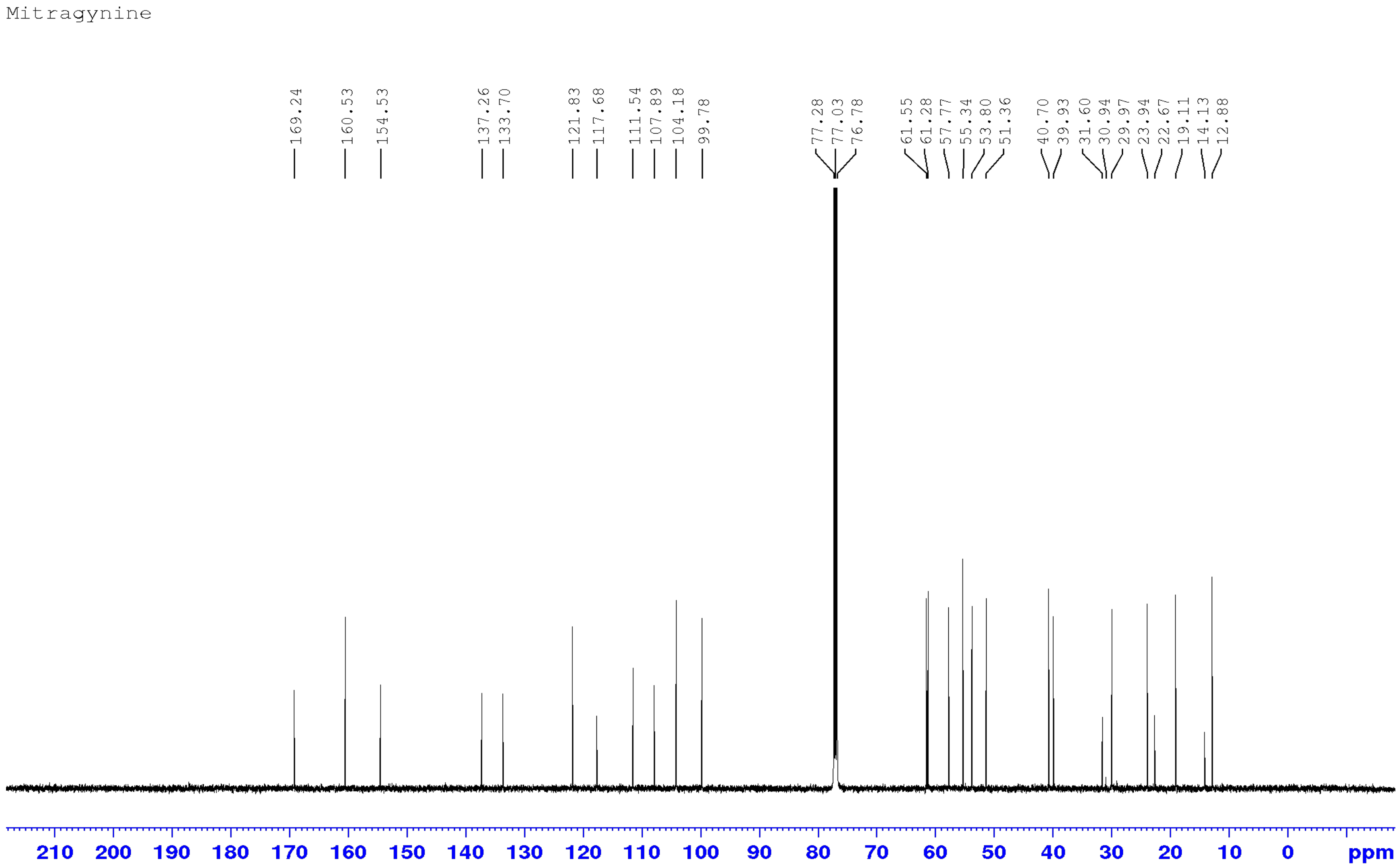


**Supplementary figure 2**: $^{13}$C NMR spectrum of mitragynine (**1**) (CDCL$_3$; 125 MHz).

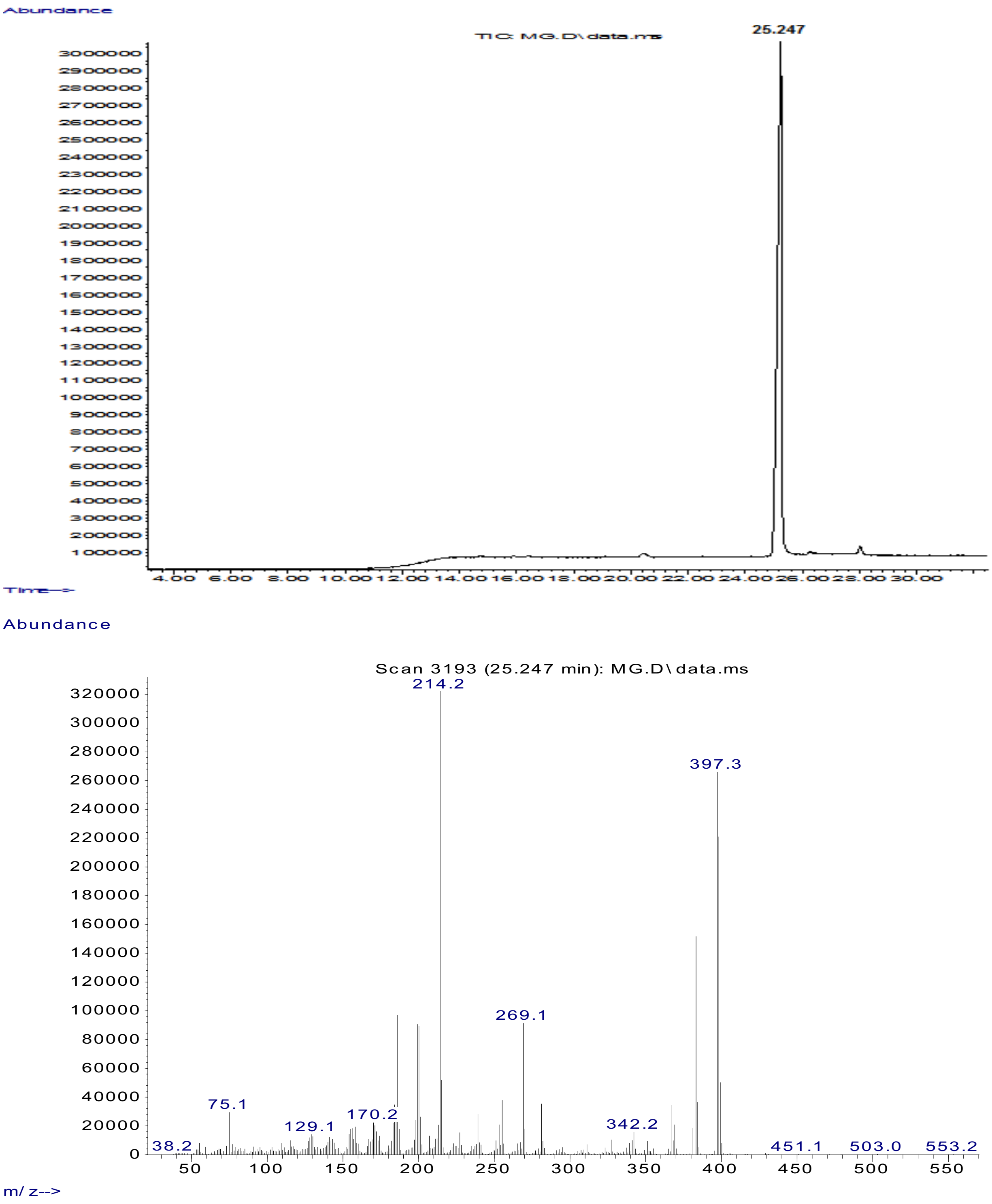


**Supplementary figure 3**: Total ion chromatogram (GC) and corresponding EI-MS spectrum of mitragynine (**1**).

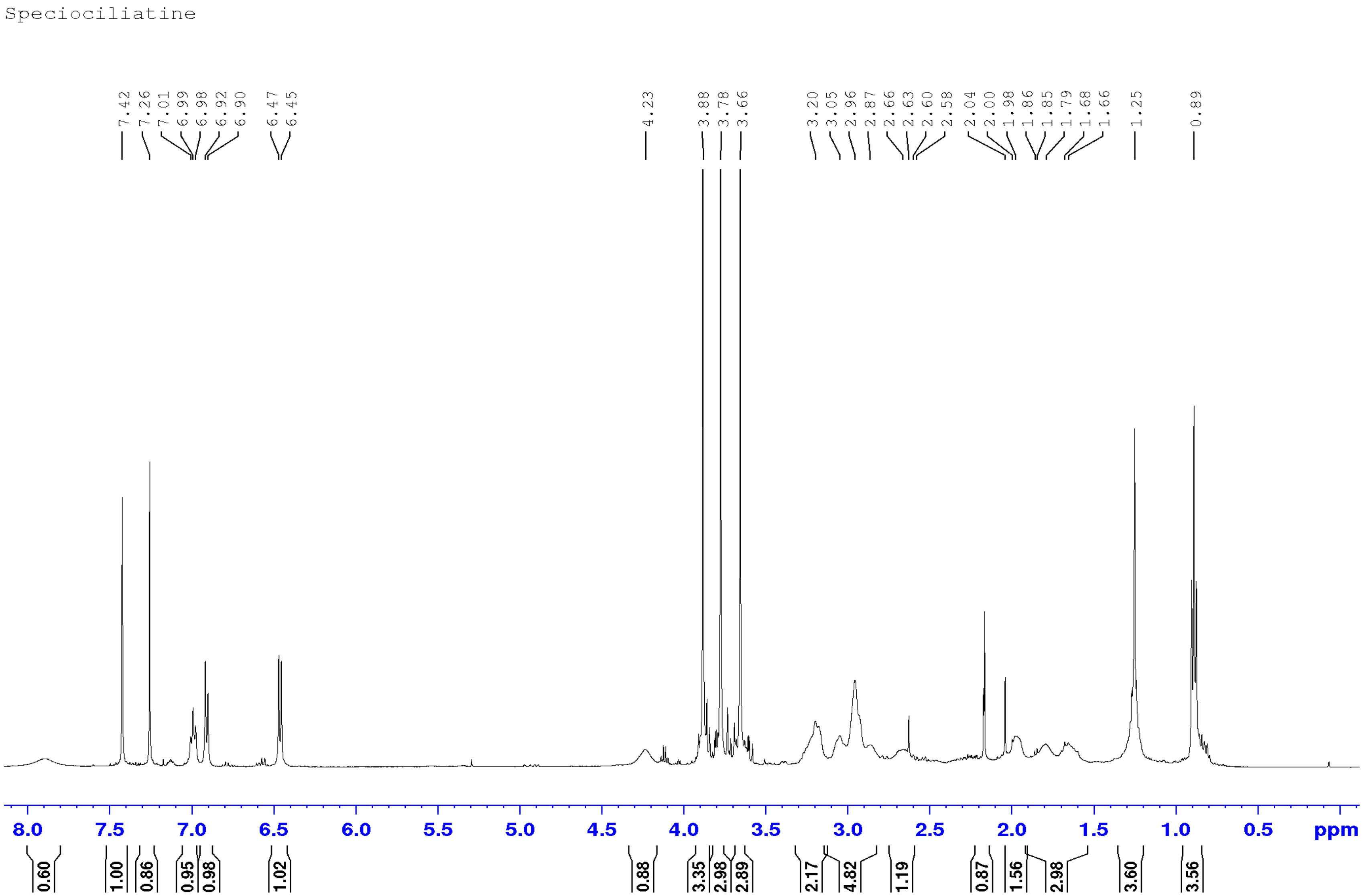


**Supplementary figure 4**: [1]H NMR spectrum of speciociliatine (**2**) ($CDCL_3$, 500 MHz).

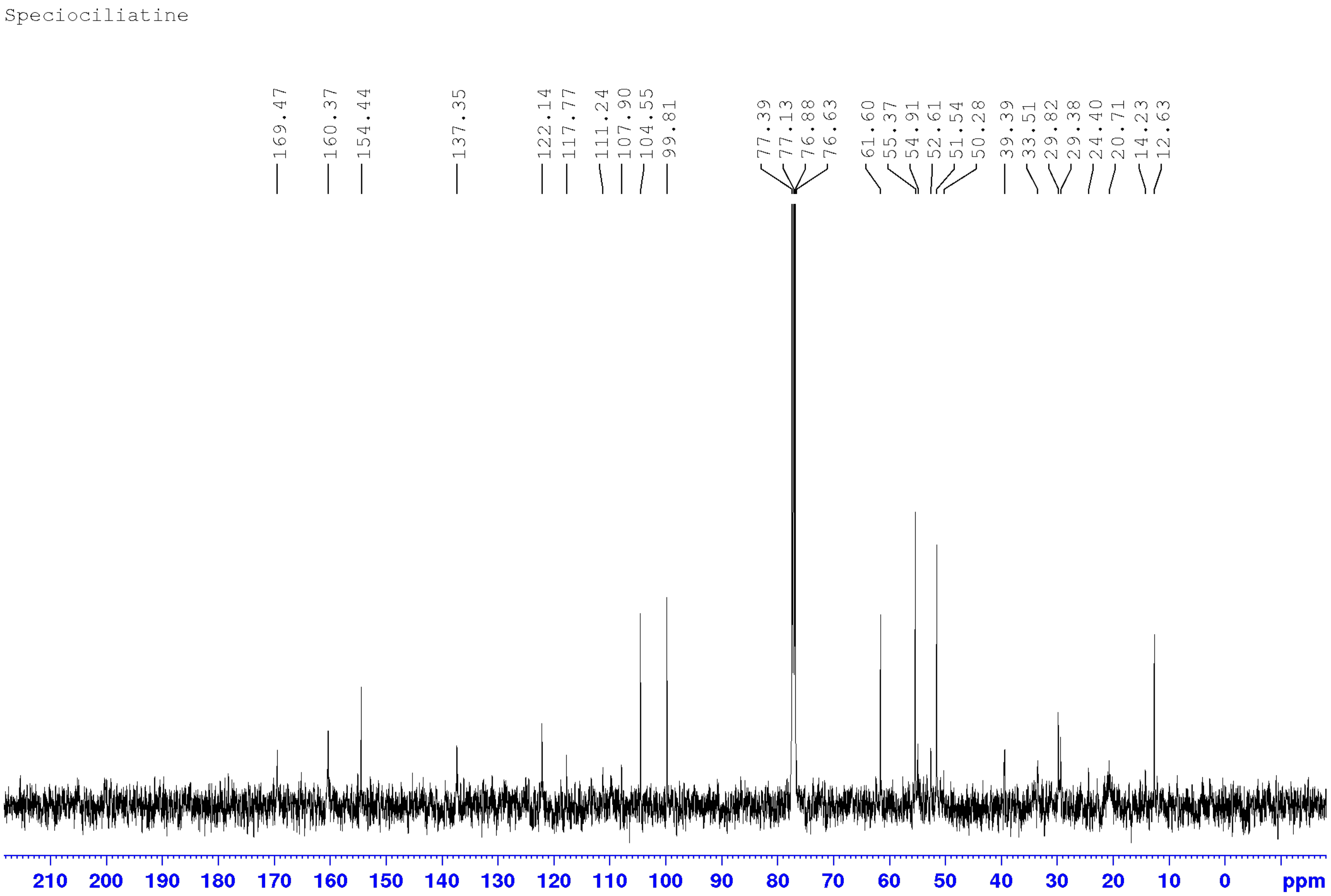


**Supplementary figure 5**: $^{13}C$ NMR spectrum of speciociliatine (**2**) ($CDCL_3$; 125 MHz).

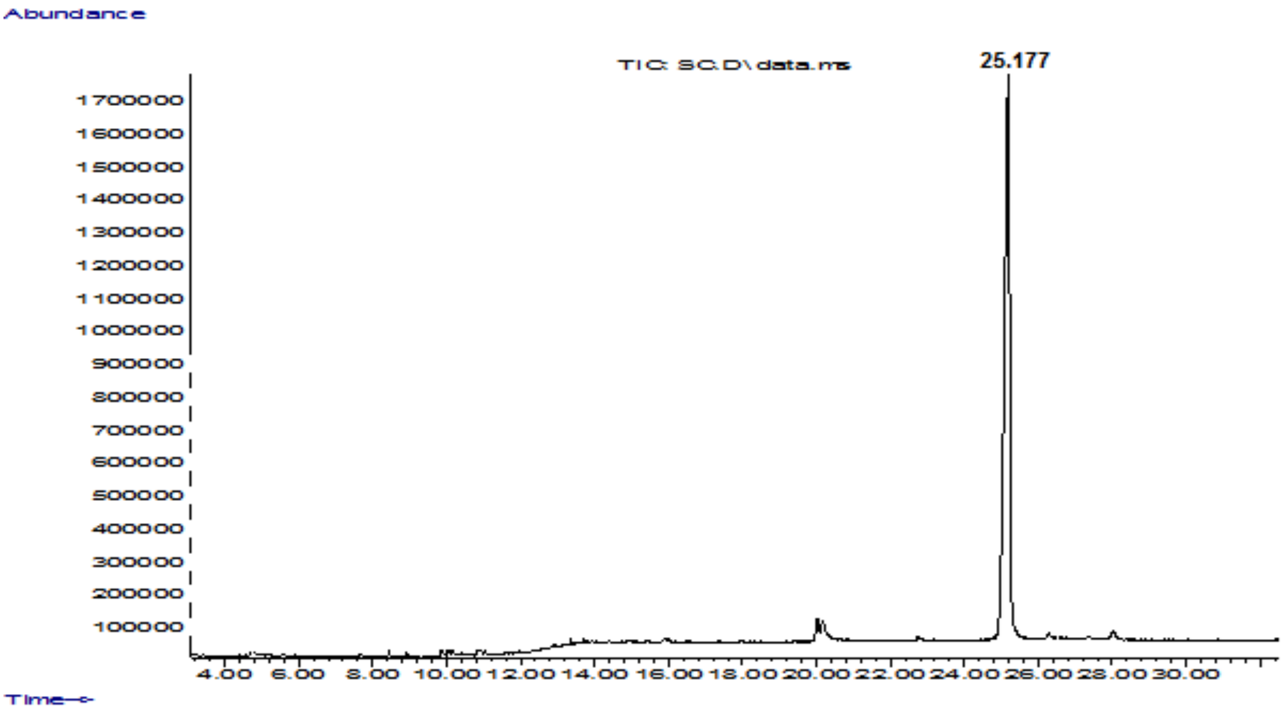


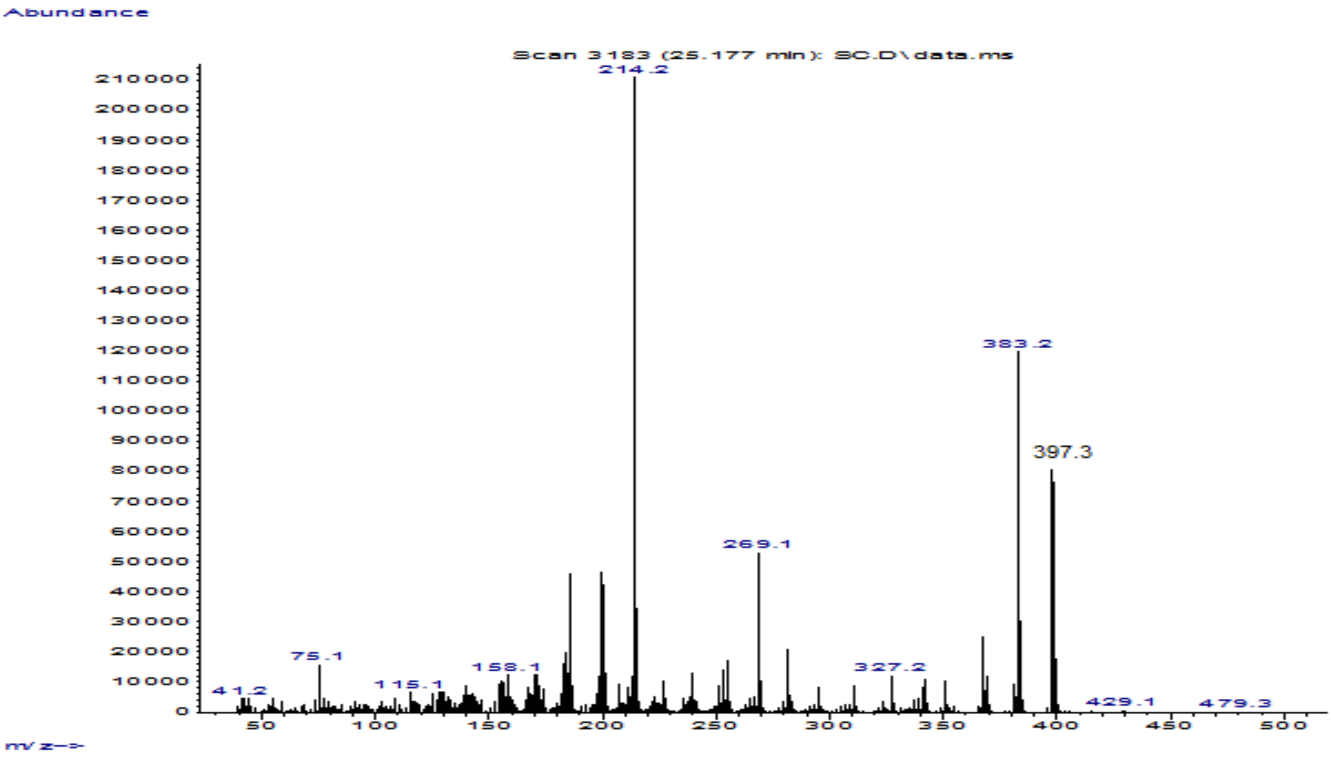


**Supplementary figure 6**: Total ion chromatogram (GC) and corresponding EI-MS spectrum of speciociliatine (**2**).

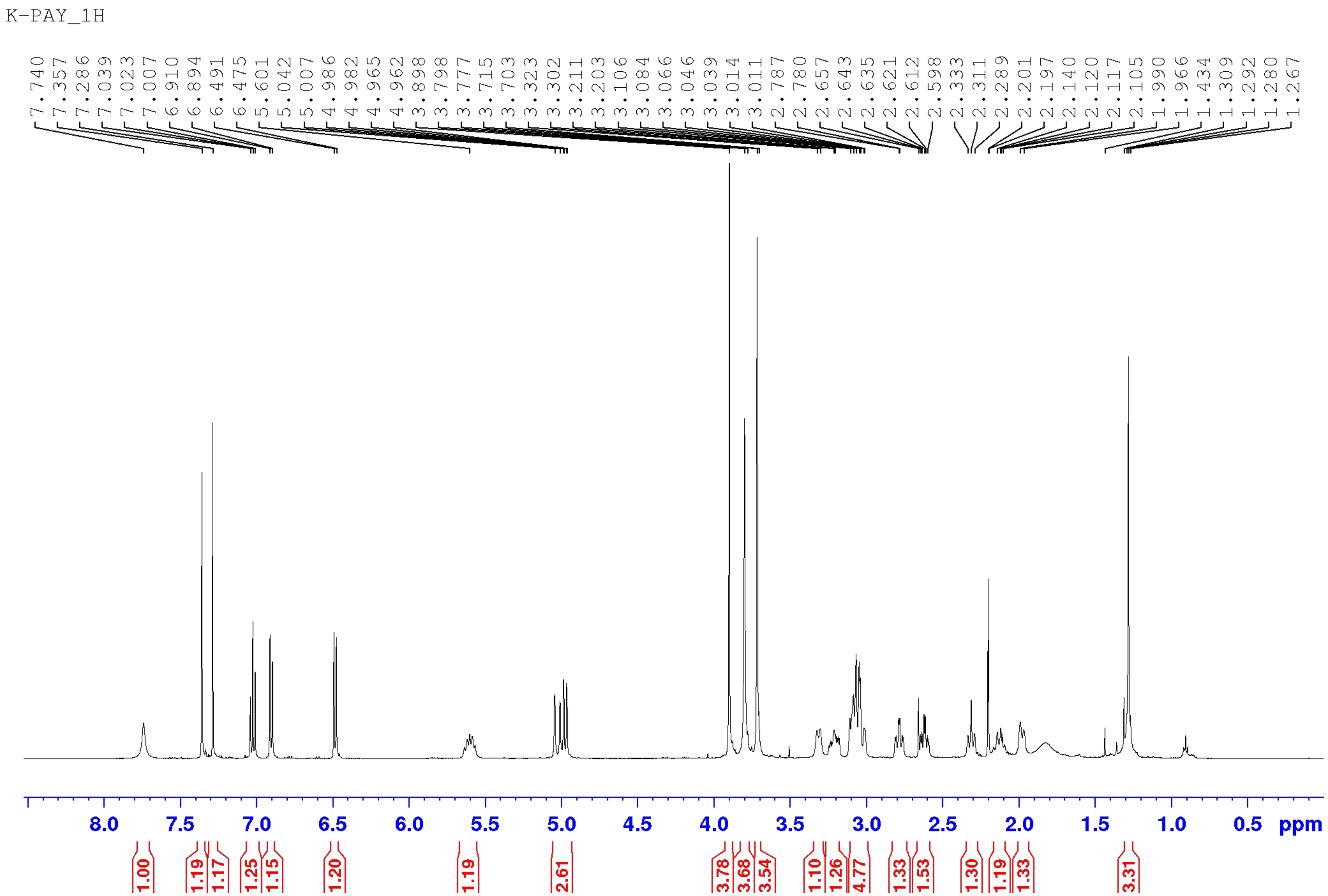


**Supplementary figure 7**: [1]H NMR spectrum of paynantheine (**3**) ($CDCL_3$, 500 MHz).

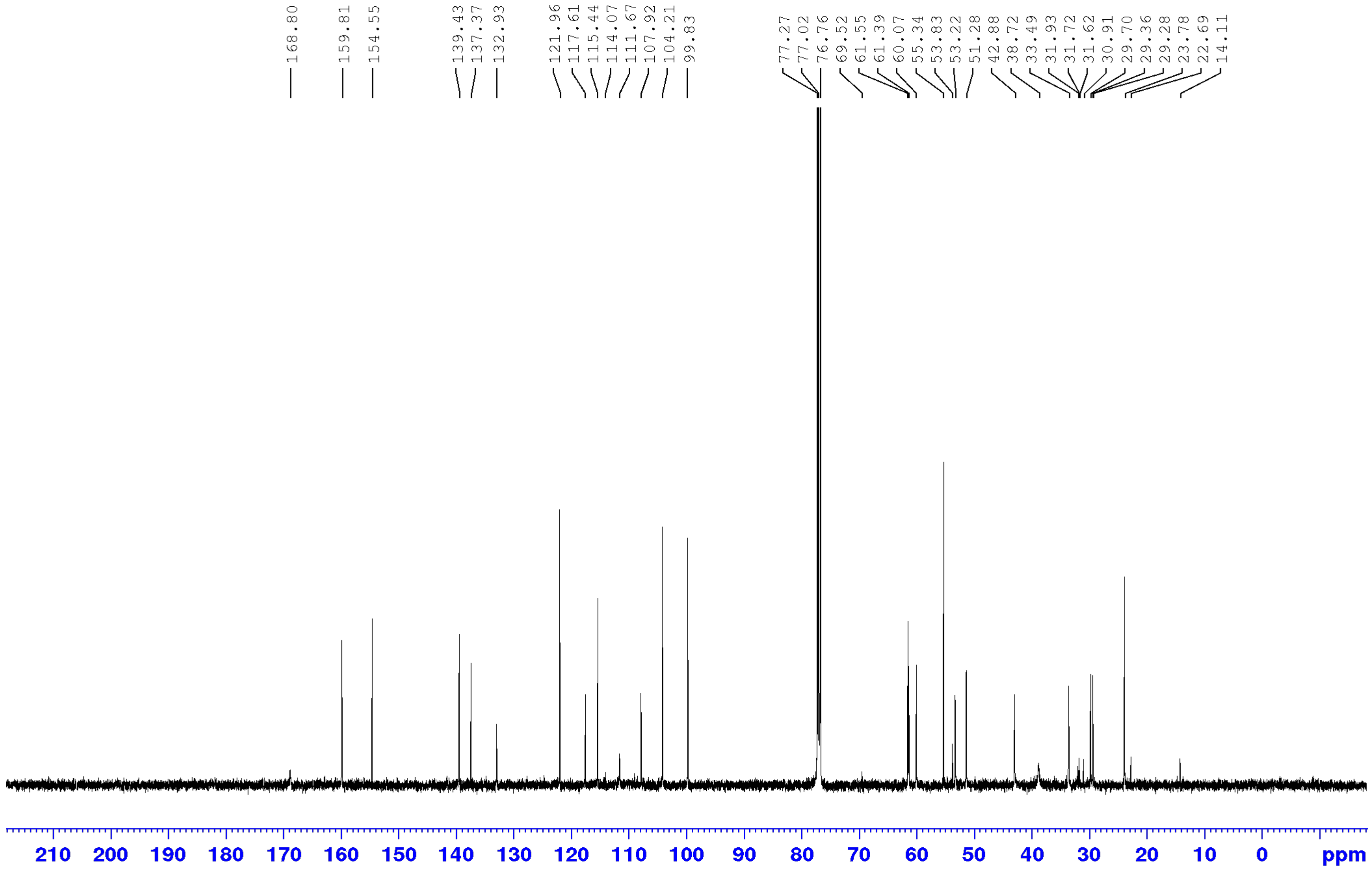


**Supplementary figure 8**: $^{13}$C NMR spectrum of paynantheine (**3**) ($CDCL_3$; 125 MHz).

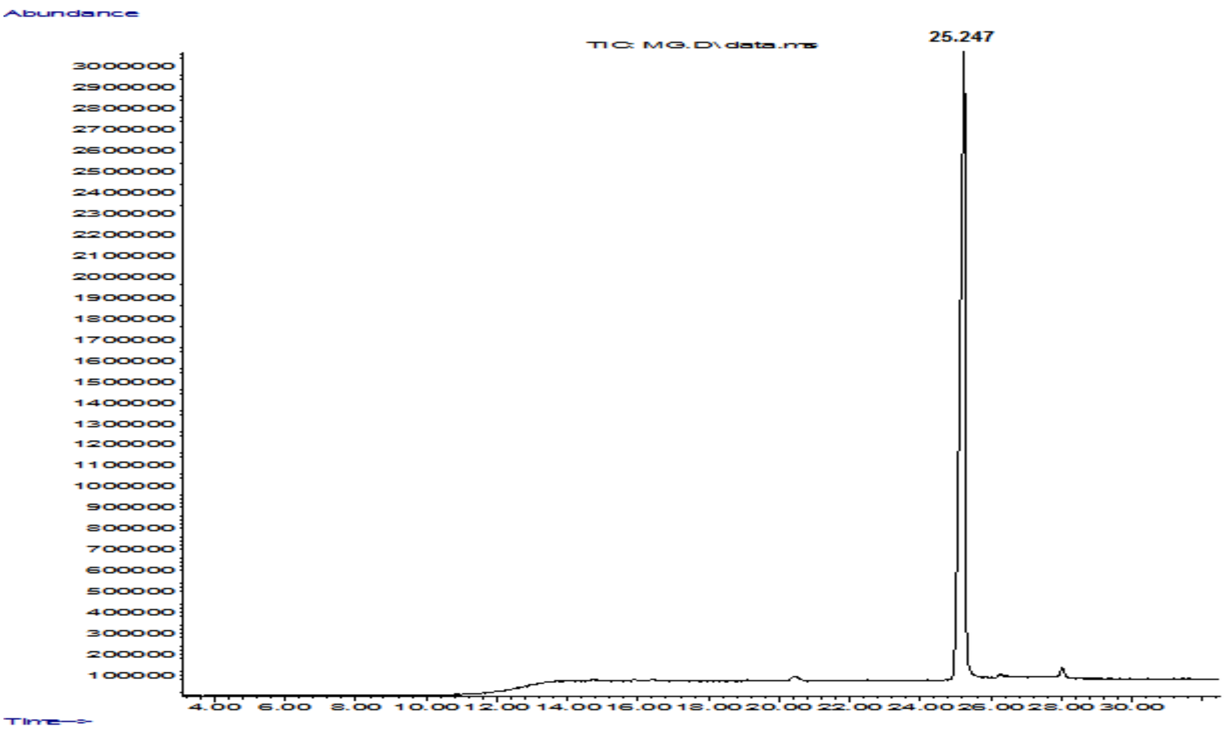


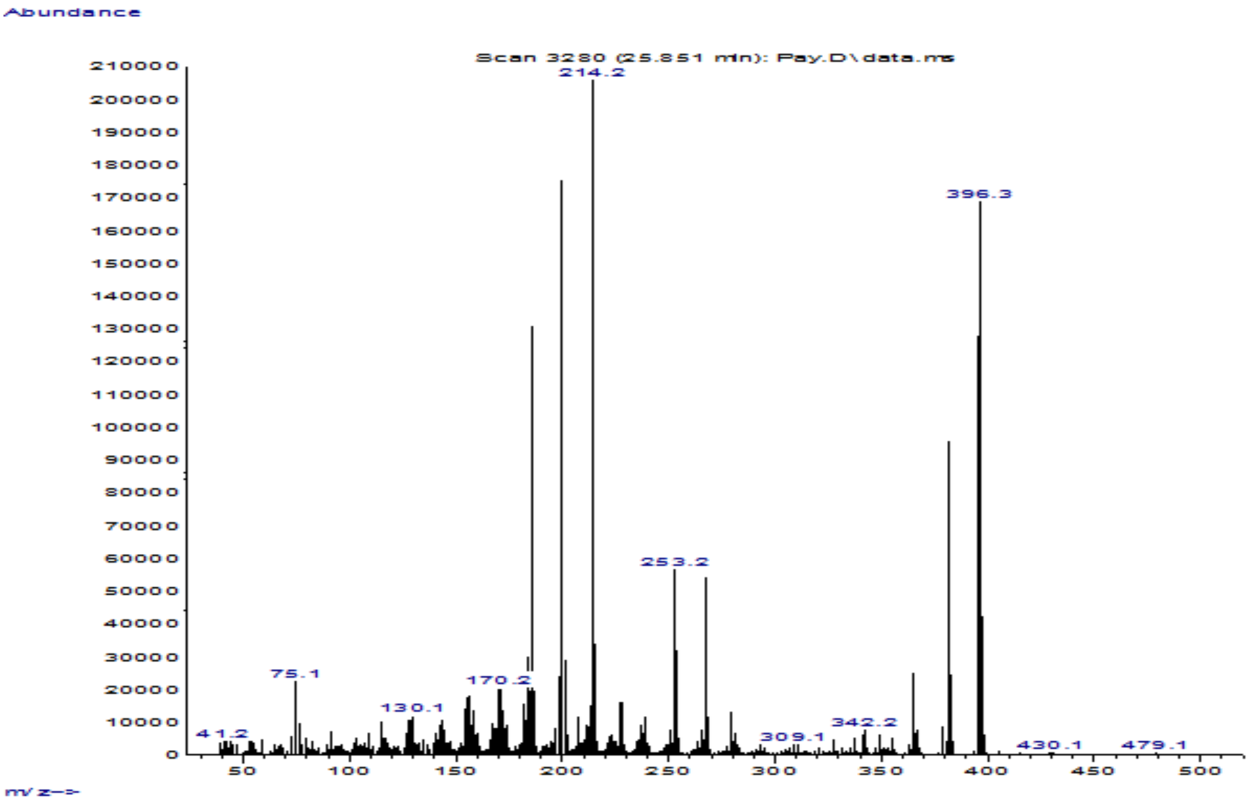


**Supplementary figure 9**: Total ion chromatogram (GC) andcorresponding EI-MS spectrum of paynantheine (**3**).

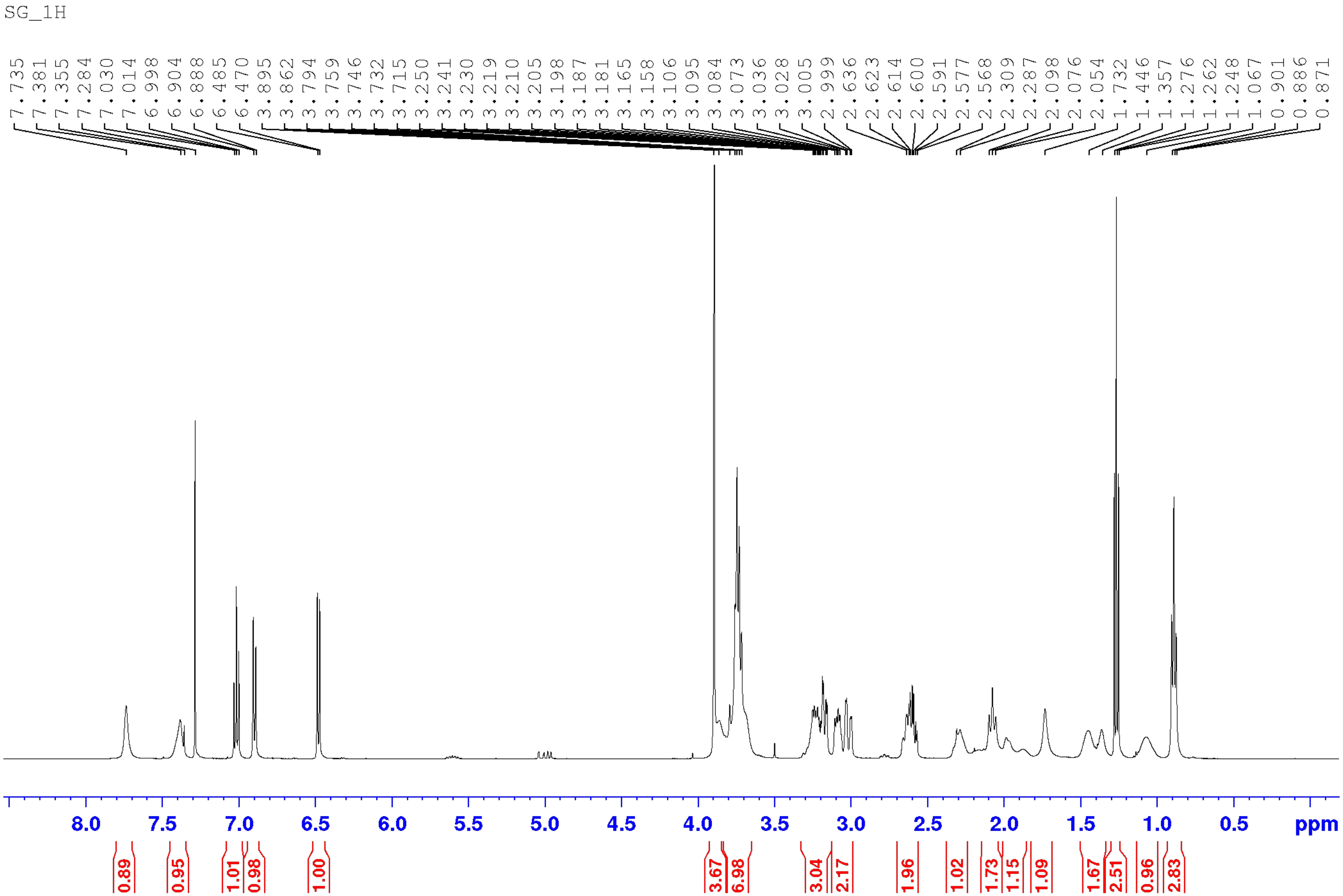


**Supplementary figure 10**: [1]H NMR spectrum of speciogynine (**4**) ($CDCL_3$, 500 MHz).

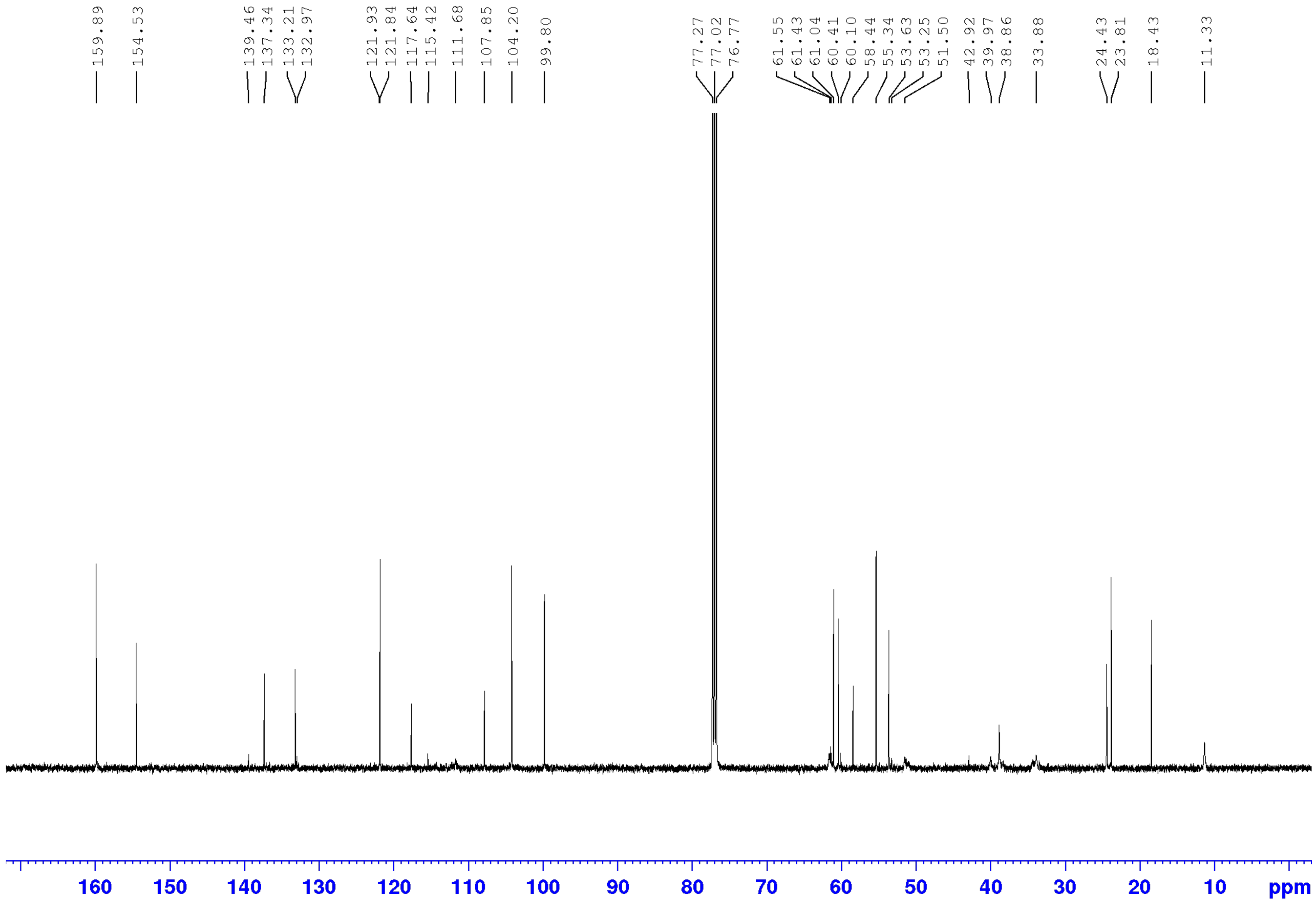


**Supplementary figure 11:** $^{13}C$ NMR spectrum of speciogynine (**4**) (CDCL$_3$; 125 MHz).

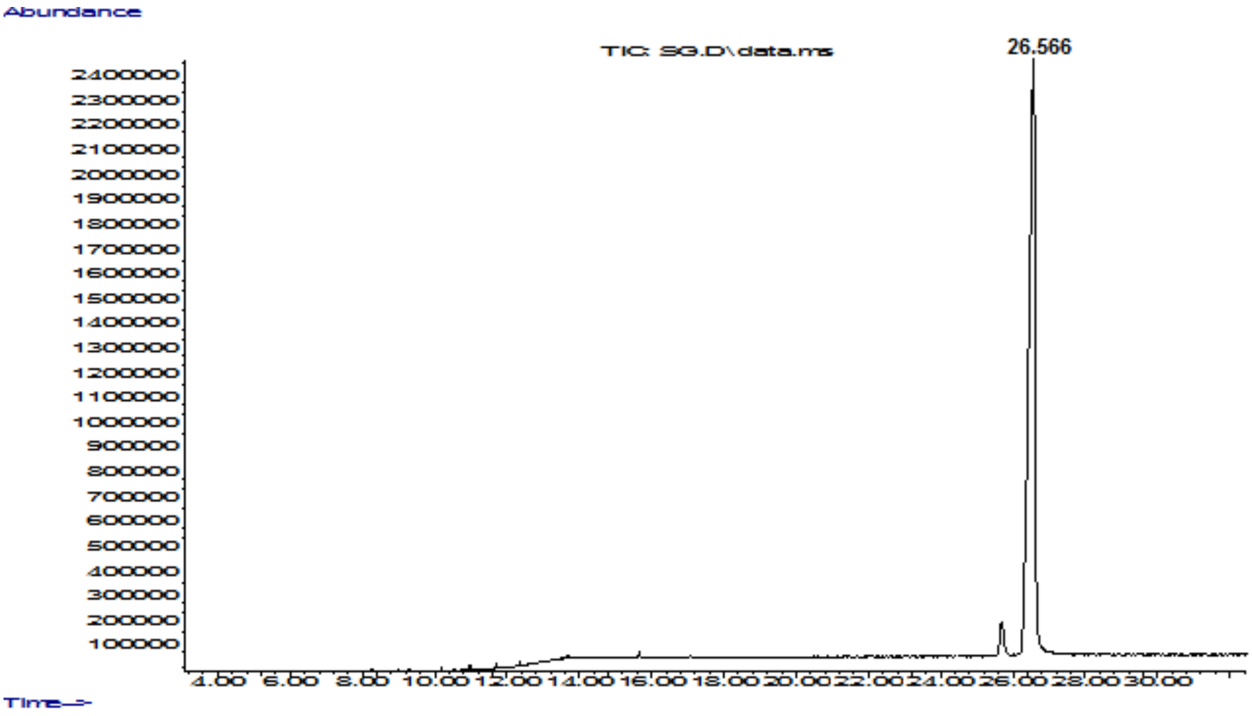


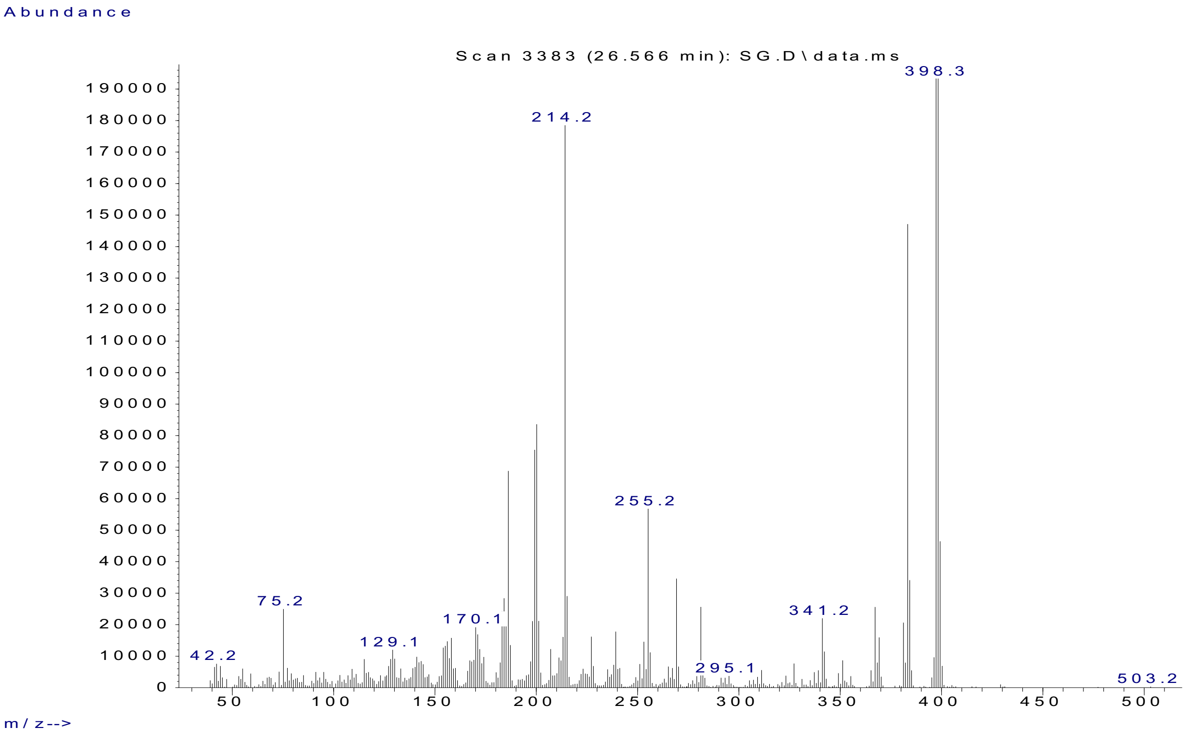


**Supplementary figure 12**: Total ion chromatogram (GC) and corresponding EI-MS spectrum of speciogynine (**4**).

| No. Peak | Retention time (mins) | Area | Area Sum % | Compound |
|---|---|---|---|---|
| 1 | 6.845 | 43.9 | 1.077 | - |
| 2 | 7.776 | 4008.1 | **98.287** | **mitragynine** |
| 3 | 17.447 | 25.9 | 0.635 | - |

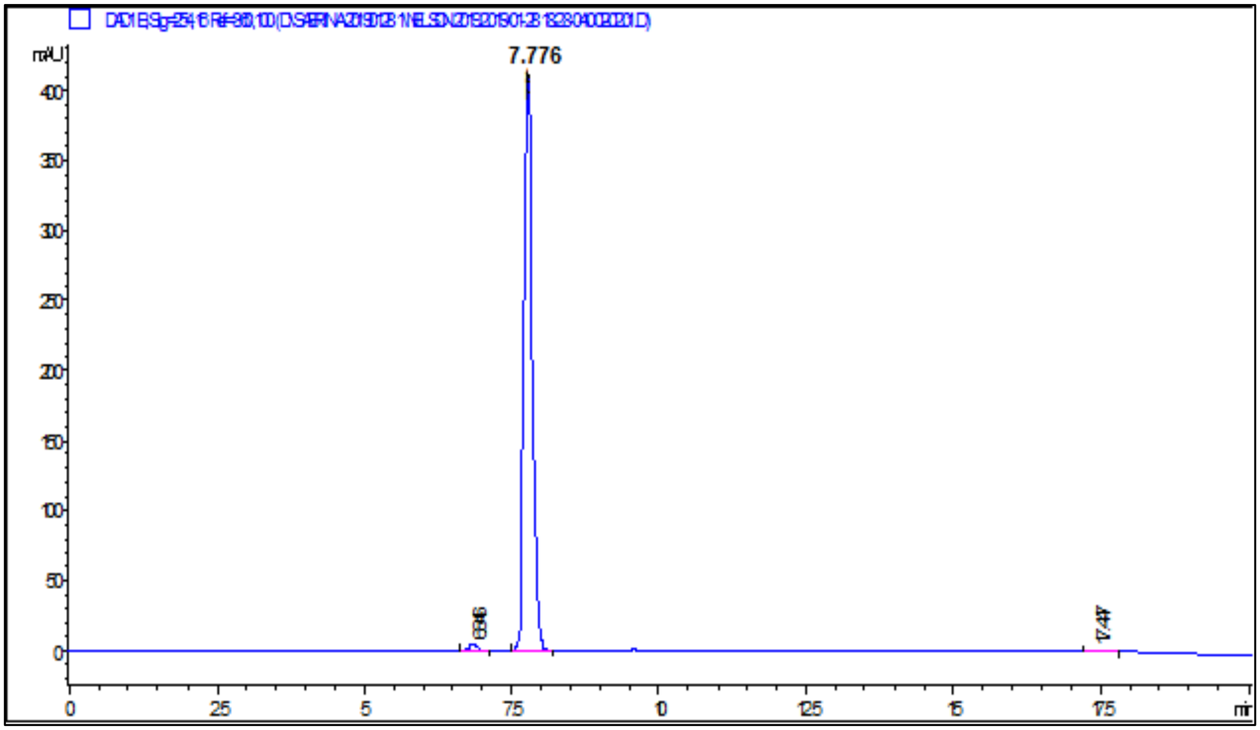


**Supplementary figure 13:** HPLC chromatogram and purity of mitragynine (**1**) at 254 nm.

| No. Peak | Retention time (RT; min) | Area | Area Sum % | Compound |
|---|---|---|---|---|
| 1 | 8.073 | 5.8 | 4.907 | - |
| 2 | 8.864 | 1062.4 | **95.093** | **Speciociliatine** |

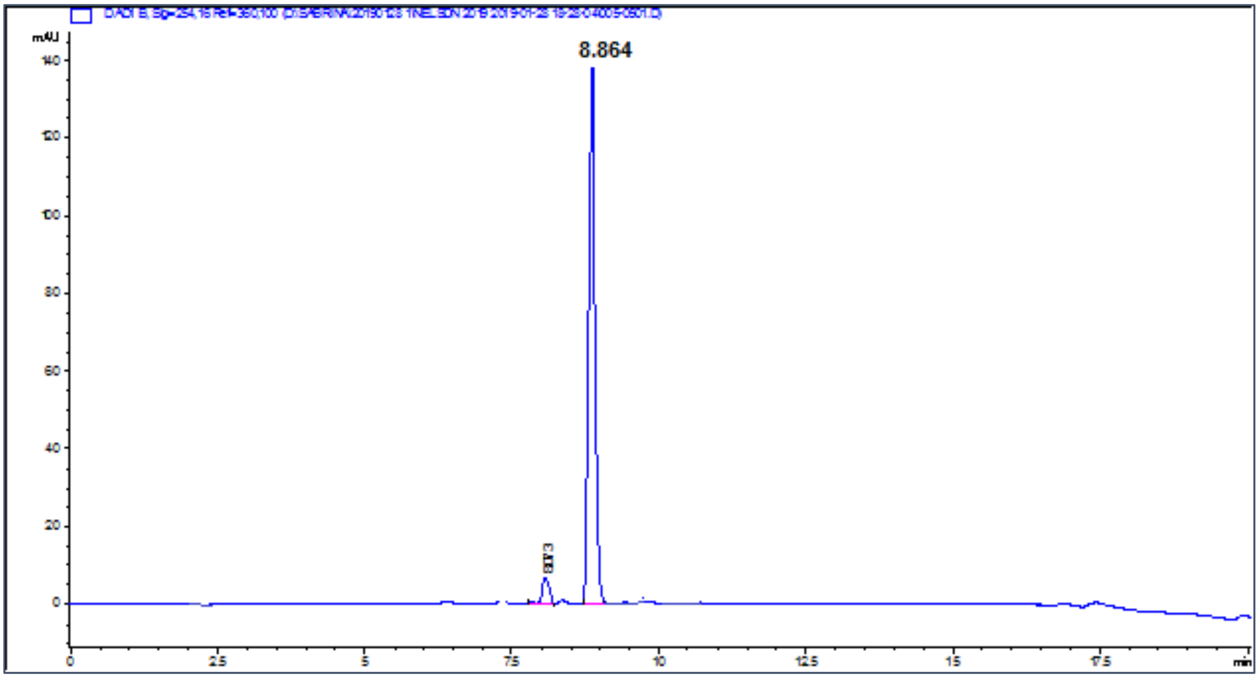


**Supplementary figure 14:** HPLC-PDA chromatogram and purity of speciociliatine (**2**) at 254 nm.

| No. Peak | Retention time (min) | Area | Area Sum % | Compound |
|---|---|---|---|---|
| 1 | 2.165 | 78.3 | 4.234 | - |
| 2 | 7.84 | 1770.5 | **95.766** | **Paynantheine** |

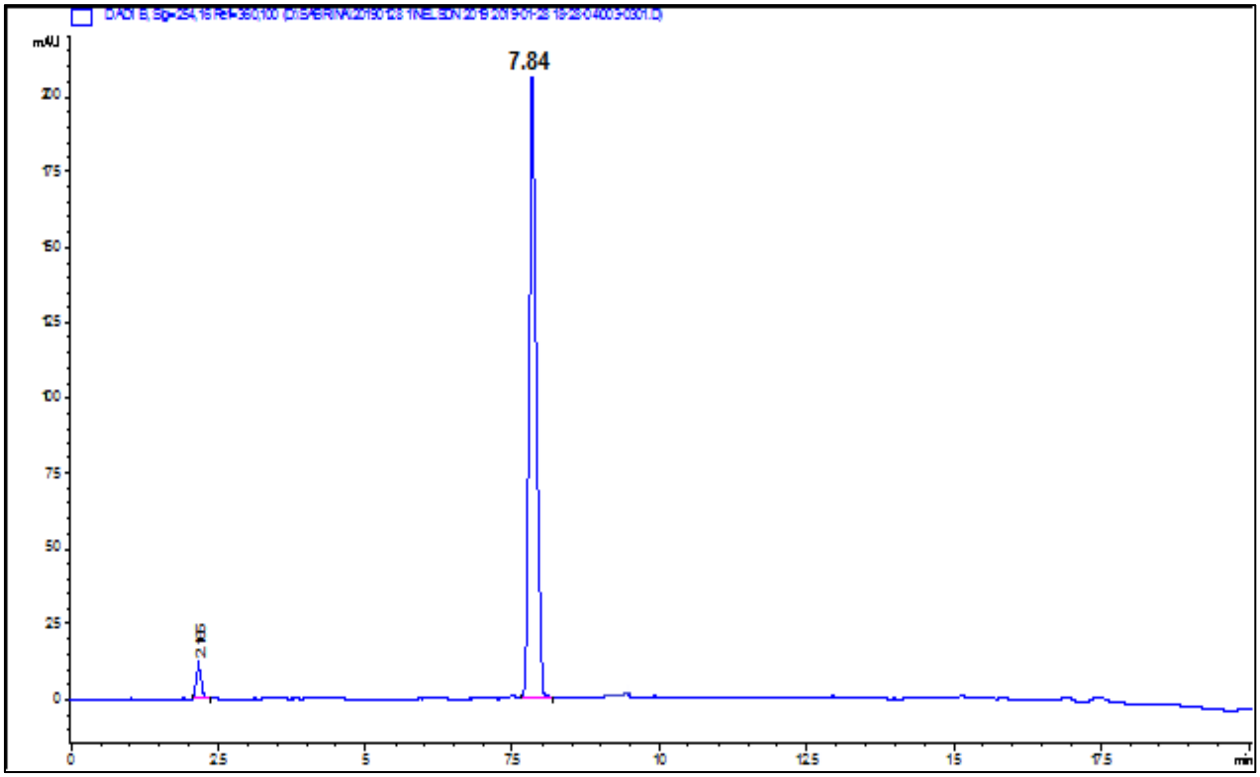


**Supplementary figure 15**: HPLC-PDA chromatogram and purity of paynantheine (**3**) at 254 nm.

| No. Peak | Time | Area | Area Sum % | Compound |
|---|---|---|---|---|
| 1 | 7.877 | 100.3 | 5.4 | - |
| 2 | 8.385 | 1778.8 | 94.6 | Speciogynine |

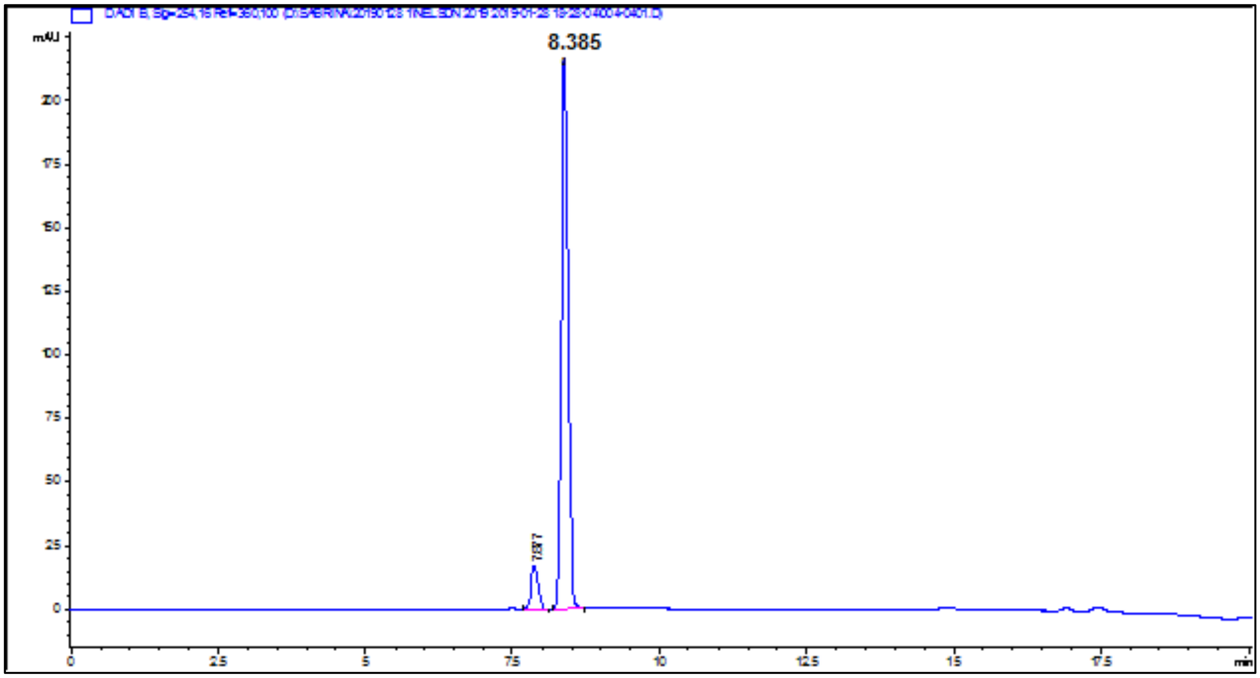


**Supplementary figure 16**: HPLC-PDA chromatogram and purity of speciogynine (**4**) at 254 nm.

**Supplementary Table 1:** $IC_{50}$ values of single agent treatment of the *M. speciosa* alkaloids on HK-1 and C666-1 cell lines.

| Cell Line | Drugs | $IC_{50}$ ± SEM (µM) |
|---|---|---|
| **HK-1** | Mitragynine | 30.61 ± 0.93 |
| | Speciociliatine | >32 |
| | Paynantheine | >32 |
| | Speciogynine | >32 |
| **C666-1** | Mitragynine | >32 |
| | Speciociliatine | >32 |
| | Paynantheine | >32 |
| | Speciogynine | >32 |

**Supplementary Table 2:** $IC_{50}$ values of combinations of *M. speciosa* alkaloids on NPC cell line HK-1.

| Cell Line | Compounds | $IC_{50}$ ± SEM (µM) | Drug Interactions |
|---|---|---|---|
| **HK-1** | Mitragynine + Speciociliatine (1:1 drug concentration ratio) | 26.10 ± 1.21 | Antagonism |
| | Mitragynine + 10 µM Speciociliatine | >30 | Slight Antagonism |
| | Speciociliatine + 10 µM Mitragynine | >30 | Nearly additive |

**Supplementary table 3: IC$_{50}$ values of the NPC cell lines to single agent treatment of cisplatin and the mitragyna alkaloids**

| Cell line | IC$_{50}$ (µM ± SEM) | | | |
|---|---|---|---|---|
| | | Mitragyna alkaloids | | |
| | Cisplatin | Mitragynine | Speciociliatine | Paynanthiene |
| HK-1 | 9.7 ± 0.4 | 30.6 ± 1.0 | > 32 | > 32 |
| C666-1 | 13.9 ± 1.0 | > 32 | > 32 | > 32 |
